# The tapeworm interactome: inferring confidence scored protein-protein interactions from the proteome of *Hymenolepis microstoma*

**DOI:** 10.1101/668988

**Authors:** Katherine James, Peter D. Olson

## Abstract

Reference genome and transcriptome assemblies of helminths have reached a level of completion whereby secondary analyses that rely on accurate gene estimation or syntenic relationships can be now conducted with a high level of confidence. Recent public release of the v.3 assembly of the mouse bile-duct tapeworm, *Hymenolepis microstoma*, provides chromosome-level characterisation of the genome and a stabilised set of protein coding gene models underpinned by both bioinformatic and empirical data. However, interactome data have not been produced. Conserved protein-protein interactions in other organisms, termed interologs, can be used to transfer interactions between species, allowing systems-level analysis in non-model organisms. Here, we describe a probabilistic, integrated network of interologs for the *H. microstoma* proteome, based on conserved protein interactions found in eukaryote model species. Almost a third of the 10,139 gene models in the v.3 assembly could be assigned interaction data and assessment of the resulting network indicates that topologically-important proteins are related to essential cellular pathways, and that the network clusters into biologically meaningful components. Moreover, network parameters are similar to those of single-species interaction networks that we constructed in the same way for *S. cerevisiae, C. elegans* and *H. sapiens*, demonstrating that information-rich, system-level analyses can be conducted even on species separated by a large phylogenetic distance from the major model organisms from which most protein interaction evidence is based. Using the interolog network, we then focused on sub-networks of interactions assigned to discrete suites of genes of interest, including signalling components and transcription factors, germline ‘multipotency’ genes, and differentially-expressed genes between larval and adult worms. These analyses not only showed an expected bias toward highly-conserved proteins, such as components of intracellular signal transduction, but in some cases predicted interactions with transcription factors that aid in identifying their target genes. With the completion of key helminth genomes, such systems level analyses can provide an important predictive framework to guide basic and applied research on helminths and will become increasingly informative as protein-protein interaction data accumulate.

## 1 Introduction

Genomic resources for parasitic flatworms and other helminths have increased substantially over the last decade. Reference genomes of key species have undergone multiple iterations of improvement, employing new sequencing and algorithmic advances to produce more contiguous assemblies and reliable estimates of coding regions and other features [Protasio et al., 2012]. At the same time, the diversity of helminth species with draft genomes continues to expand [International Helminth Genomes Consortium, 2019], enabling work on a broader range of species and more informative comparative analyses. Among flatworms, the human bloodfluke *Schistosoma mansoni* and the tapeworms *Echinococcus multilocularis* and *Hymenolepis microstoma* are now supported by near complete, chromosome-level assemblies, providing more comprehensive and stable gene model estimates and syntenic relationships, as well as allowing the higher order architecture of their genomes to begin to be investigated. The unusually high level of completeness and quality of these assemblies makes them valuable not only for investigating these taxa, but also as models of the superphylum Lophotrochozoa which remains significantly under-represented in most areas of biological research.

*Hymenolepis microstoma*, the mouse bile-duct tapeworm, is one of three species of rodent/beetle-hosted hymenolepid tapeworms that have been used widely as laboratory models, as their entire life cycles can be passaged using hosts that are themselves model organisms [Cunningham and Olson, 2010]. A draft genome was published in 2013 [Tsai et al., 2013] and was followed in 2015 by the public release of an up-dated assembly (v.2) based on additional Illumina data, as described by Olson and colleagues [Olson et al., 2018]. This assembly was used to investigate differentially-expressed genes among different life cycle stages and regions of the adult, strobilar worm [Olson et al., 2018], and for characterisation of the microRNA complement [Macchiaroli et al., 2019]. In 2018 long-read sequence and optical mapping data were added and all available genome data re-assembled, resulting in a complete assembly consisting of six scaffolds that correspond to their six haploid chromosomes [Hossain and Jones, 1963]; any missing data that remain are likely to represent collapsed repeats rather than unique, non-repetitive sequence (Olson, in preparation). The 169 Mb v.3 assembly, including 10,139 gene models and an additional 1,290 splice variants, as well as RNA-seq data sets, is publicly available via WormBase Parasite^1^ [Howe et al., 2017]. Thus, with the basic assembly and annotation of these inaugural helminth sequencing projects now effectively complete, we can begin to undertake systems-level analyses in parasitic flatworms for the first time.

Protein-protein interactions in cellular networks are known to be highly conserved [Sharan and Ideker, 2006, von Mering et al., 2003]. Evidence suggests that a simple set of rules characterizes all protein interaction networks [Barabsi and Oltvai, 2004], with network hubs (highly-connected proteins) being conserved and essential [He and Zhang, 2006, Jeong et al., 2001, Brown and Jurisica, 2007], and having slower evolutionary rates [Fraser et al., 2002] and significant sequence conservation [Nguyen Ba et al., 2012]. Despite high-throughput interaction data having estimated false positive and negative rates as high as 90% and 50% [von Mering et al., 2002], respectively, the conservation of ‘hub’ proteins and their interactions remains detectable within eukaryotic species [Wuchty et al., 2006], and even between eukaryotes and prokaryotes [Kelley et al., 2003]. Conserved interactions, termed ‘interologs’, can therefore be transferred between species [Matthews et al., 2001, Castillo-Lara and Abril, 2018, Gu et al., 2011, Lin et al., 2011, Yellaboina et al., 2008, Bhardwaj et al., 2016, Titz et al., 2008, Musungu et al., 2015], allowing systems-level analysis in organisms lacking empirical interaction data.

Here we produced a probabilistic, integrated network of interologs for *H. microstoma* using physical interaction data from sixteen different species obtained from the BioGRID database [Stark et al., 2006]. Probabilistic networks are more powerful than unweighted networks as they are annotated with a level of confidence in the evidence for each interaction by comparison with a benchmark ‘gold standard’ comprising a set of interactions believed, with high confidence, to be true interactions [Lee et al., 2004]. This benchmarking reduces noise from high-throughput data sets, produces consistent integration of interactions from different studies, and allows the use of thresholding and statistical algorithms that take these probabilities into account. We assessed the network by comparison of the major network parameters against networks of major model organisms produced using the same methods. We then used the network to identify highly-connected ‘hub’ proteins, network clusters and interacting partners of genes of interest, including signalling components, transcription factors and germline ‘multipotency’ genes, as well as genes differentially expressed between life stages. All data in our interaction network can be readily interrogated using Cytoscape [Shannon et al., 2003], enabling users to explore predicted protein interactions for their own genes of interest. We expect that such network analyses will become an increasingly valuable resource for hypothesis generation, including predicting protein choke points for mitigating parasitism, determining processes shared between parasitic and free-living flatworms [Olson, 2008] and between flatworms and ourselves [Castillo-Lara and Abril, 2018].

## 2 Materials and Methods

### 2.1 Network Integration

Networks were derived using a four-stage scoring, filtering, integration and thresholding method (Figure 1A). Interaction data were downloaded from BioGRID (ver. 164). BioGRID is a comprehensive and highly-curated resource for functional association data [Stark et al., 2006]. The database stores interactions of 28 different types, including both physical interactions (17 types), for instance from affinity capture and yeast two hybrid studies, and genetic interaction evidence (11 types), such as dosage or synthetic growth defects. We filtered the data to remove non-eukaryotic and non-physical interaction types. Data were split into individual data sets by study and species, with low-throughput (LTP) studies (*<* 200 interactions) grouped by experimental type (Tables S1), while high-throughput (HTP) studies (*>*= 200 interactions) were treated as separate datasets (Table S2). BioSystems pathways (downloaded 20^*th*^ February 2019) were used as the gold standard for confidence scoring. Confidence scores were calculated using the methods developed by Lee and colleagues [Lee et al., 2004], that calculates a log-likelihood score for each data set (1):

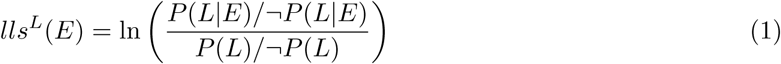

where, *P* (*L|E*) and *¬P* (*L|E*) represent the frequencies of linkages *L* observed in dataset *E* between genes annotated to the same and differing BioSystems pathways, respectively, and, *P* (*L*) and *¬P* (*L*) represent the prior expectation of linkages between genes in the same and differing BioSystems pathways, respectively. Since interaction and gold standard data for some species were very sparse, a baseline count of one was used in all cases to ensure minimal loss of these datasets. A score greater than zero indicated that a dataset links genes annotated to the same pathway; higher scores indicate greater confidence in the predicted interactions. Datasets that did not have a positive score were discarded.

**Figure 1.**
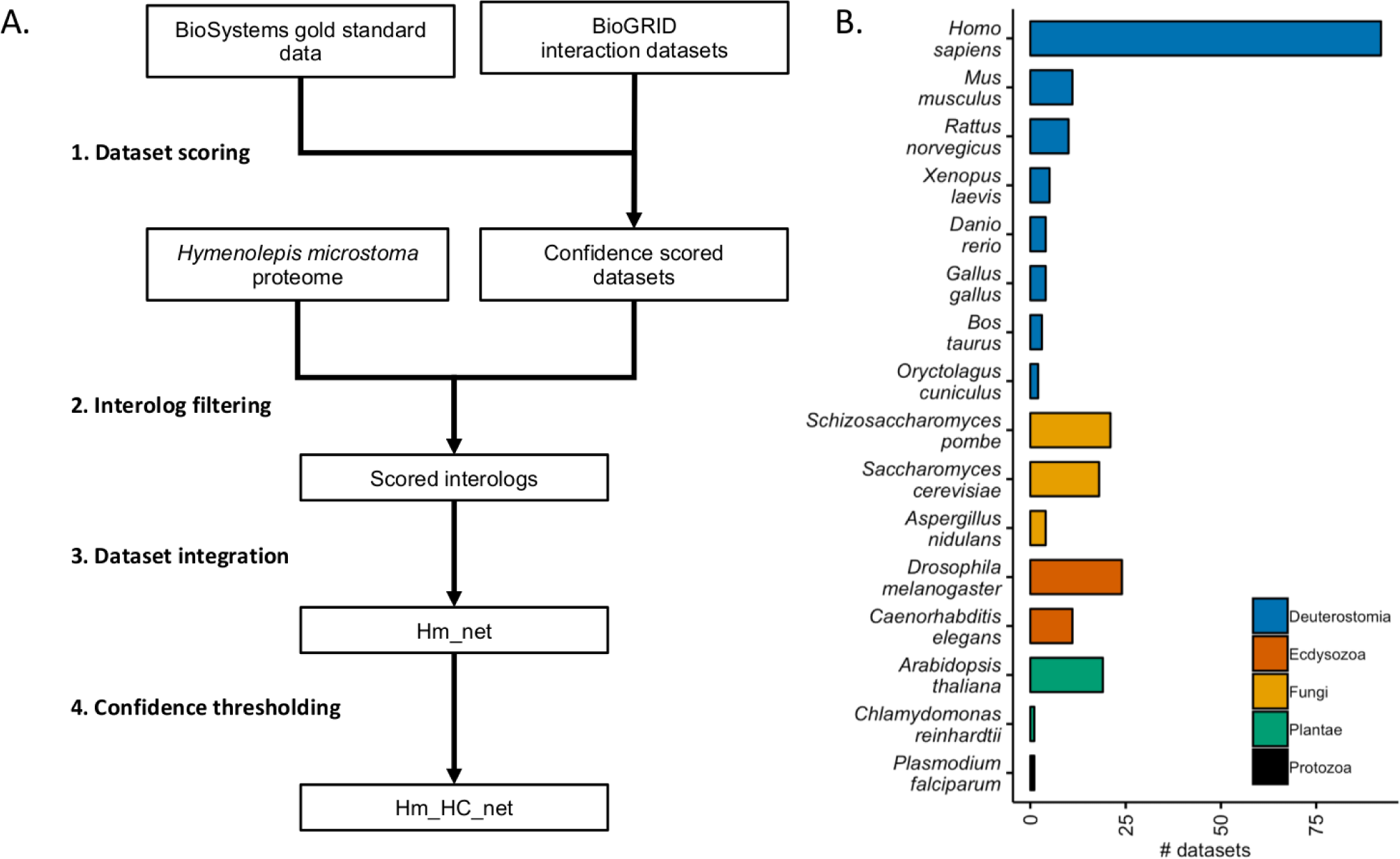
Network integration. A. The *H. microstoma* interolog network was integrated using a four-stage scoring, integration, filtering and thresholding method. First, physical interaction from the BioGRID dataset were confidence scored against gold standard BioSystems data (1). These datasets were then filtered for interactions with Blast hits to the *H. microstoma* proteome (2) before integration using a weighted sum to produce the full interolog network, Hm net (3). Finally, the network was thresholded based on interaction confidence to produce a high confidence network, Hm HC net (4). B. Data from sixteen eukaryotic species were integrated in to Hm (classification groups are based on

Orthologs of the *H. microstoma* proteome (version v.3) were identified with Blast+ (version 2.7.1) using the --gilist option to limit the search to NCBI identifiers from species in the BioGRID database (e-value *<* 0.00001), and the results filtered for the top hit to BioGRID interacting proteins in each species. Identifier mappings were obtained from the UniProt [UniProt Consortium, 2018] ftp server (downloaded 21^*st*^ February 2019). All *H. microstoma* splice variants were treated as a single protein to avoid redundant interactions. The BioGRID datasets were then filtered to retain interactions involving those proteins with orthologs (i.e. interologs), before being integrated using the Lee method [Lee et al., 2004] with a *D* -value of 1.0:

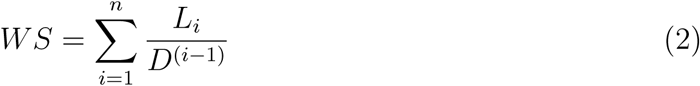

where *L*_1_ is the highest confidence score and *L*_*n*_ the lowest confidence score of a set of *n* datasets.

Networks for human, yeast and worm were derived by integration of the unfiltered data sets for that species using the same integration method.

### 2.2 Network Analysis

Network data were visualised in Cytoscape (version 3.7.1) [Shannon et al., 2003]. Network statistics and plots were produced using the NetworkAnalyser plugin (version 2.7) [Assenov et al., 2008]. Clustering was carried out using MCODE version 1.5.1 [Bader and Hogue, 2003] with a degree threshold of 3, node score threshold of 0.2 and the ‘haircut’ option.

To examine if we could predict proteins that interact with specific genes of interest (Table S3), we asked whether orthologs from four protein sets were present in the network, and where so, extracted the relevant sub-networks to examine the interologs:

1. Components of the Wnt, Notch and Hedgehog signalling pathways [Tsai et al., 2013]
2. Hox family homeobox transcription factors [Tsai et al., 2013]
3. Germline ‘multipotency’ genes [Juliano et al., 2010].
4. Differentially-expressed genes with a log2 fold-change *≥* 2 between whole, gravid adults and 5-day old larval worms [Olson et al., 2018].

Differential expression was calculated by re-analysing previously available RNA-seq data [Olson et al., 2018] in order to take advantage of the v.3 genome assembly and gene models. Briefly, raw reads were aligned to the genome using STAR [Dobin et al., 2013] v2.4.2a with the alignIntronMin 10 option, count files were produced using featureCounts v1.6.3 [Liao et al., 2014], and differential expression assessed using DESeq2 [Love et al., 2014] v1.20.0 with a *p*-value threshold of 0.00001. These results supercede those based on the v.2 genome in [Olson et al., 2018], and a full list of differentially-expressed genes among all of the RNA-seq samples generated in [Olson et al., 2018] is included in Olson (in preparation).

## 3 Results

We integrated a probabilistic network of *H. microstoma* interologs using a four-step scoring, filtering, integration and thresholding pipeline (Figure 1A; Methods). BioGRID datasets were first filtered to remove bacterial data before confidence scoring against a gold standard data set derived from the BioSystems database. A total of 528 datasets were produced (Tables S1 and S2), 428 of which had a positive confidence score (Figure S1). Blastx was then used to identify hits against the *H. microstoma* proteome within the available species. Finally, the Blast hits were mapped to the scored datasets and the dataset confidence scores integrated using a weighted sum (Methods). In total, 230 data sets from 16 species were included in the final integration step (Figure 1B), resulting in a network of 3,474 proteins (*∼* 30% of the *H. microstoma* proteome) and 20,684 interactions: Hm net (Figure 2, upper). The network scores were also filtered using a threshold to produce a high confidence sub-network of 1,494 proteins and 4,139 interactions with the highest weighted evidence: Hm HC net (Figures 2, lower and S2 A). The full tapeworm interolog network, Hm net, high confidence sub-network, Hm HC net and network annotations are provided in the Supplementary Data file (supp data.zip) in a tab-delimited format suitable for use with Cytoscape and other network analysis software.

**Figure 2.**
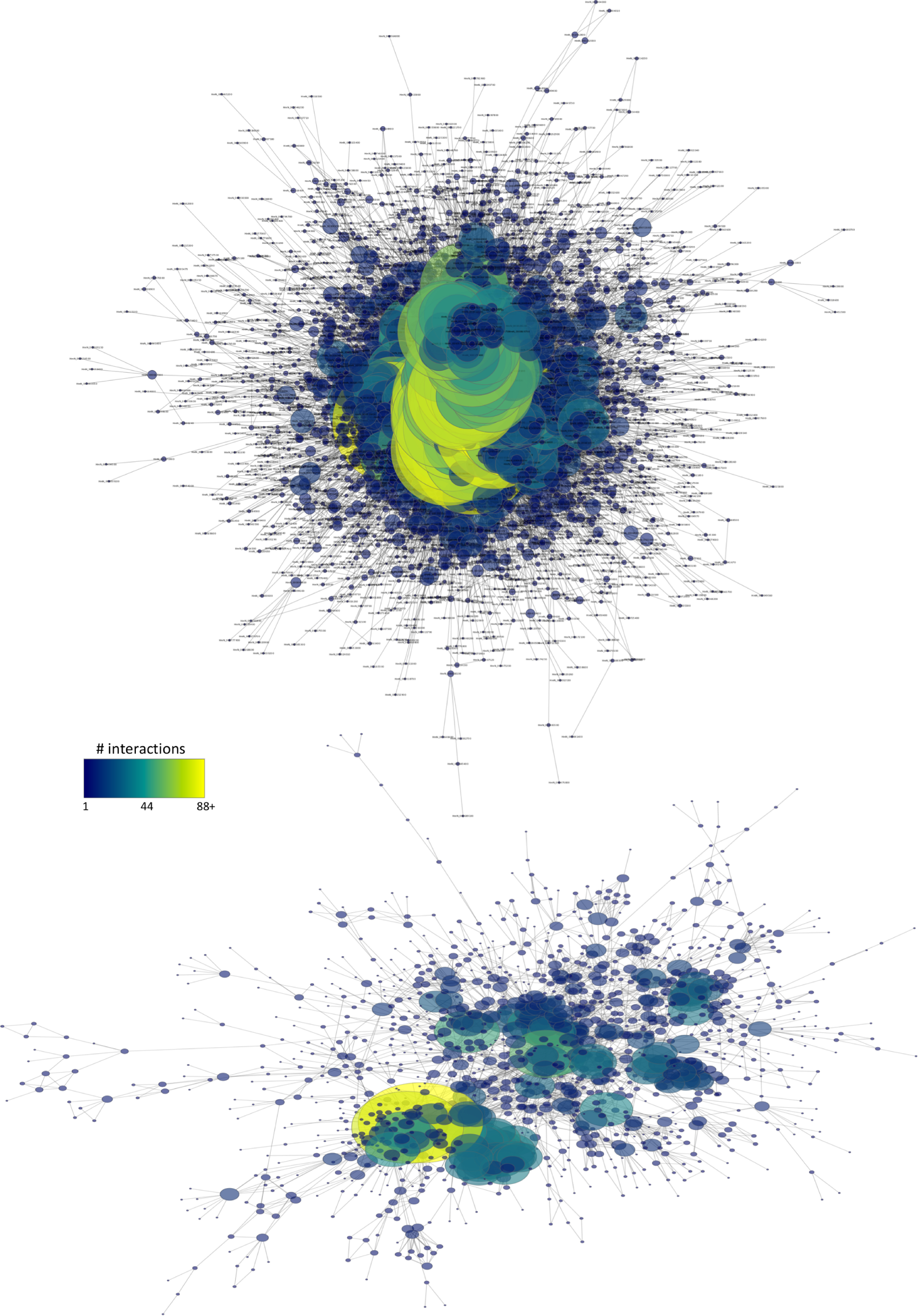
Hm_net. Hm net (upper) comprising 3408 proteins, *∼* 30% of the *H. microstoma* proteome, and 20640 interactions (largest connected component is shown here). The largest component of Hm_HC_net (lower), comprising 1260 proteins and 3995 interactions with the highest confidence scores. In both cases protein nodes are coloured and sized by number of interactions. Two large hub proteins (yellow) remain following the thresholding.

We assessed the network using a variety of network analysis techniques. Initially, we compared the network to protein-protein interaction networks from three model species: human, yeast, and *C. elegans* to determine how closely the topology of Hm net resembles networks produced directly from experimental protein-protein interaction data. We then investigated the topologically-important proteins and network clusters of Hm HC net in an exploratory manner. Finally, we used the network to ask whether it could predict interaction partners for groups of genes relating to development and to genes differential expressed between larval and adult worms, as enumerated above.

### 3.1 Hm net is topologically comparable to protein-protein interaction networks from major model organisms

Network topological parameters are often used to characterise the global properties of biological networks [Assenov et al., 2008]. We compared the topology of the *H. microstoma* interolog network to those of humans, yeast and *C. elegans*, integrated using the same probabilistic methodology, in order to asses how well Hm net resembles a real protein-protein interaction network. The human network consisted of 153 data sets (16 LTP), yeast 89 data sets (17 LTP) and *C. elegans* 16 data sets (14 LTP). The *H. sapiens* and *S. cerevisae* networks had a similar confidence score distribution to Hm net with a large proportion of interactions scoring below 2.5, while the majority of *C. elegans* scored below 1.0 (Figure S2 B).

The human network was by far the largest of the four networks, reflecting the larger proteome and multiple tissue types (Table 1). In contrast, the yeast network was the most dense having the smallest diameter (i.e. minimum number of links that separate the two most distant proteins in a network) and had a single connected component of interactions, reflecting its single cell type and large proportion of high-throughput data sets. Although containing fewer proteins, Hm net is similar to the other networks in terms of its overall topology. Hm net, has the highest clustering coefficient of the four, likely due to the multiple sources of interolog evidence resulting in denser connectivity between related proteins. By contrast, the *C. elegans* network is smaller and more dispersed with a larger diameter and characteristic path length.

**Table 1.** Network topology. The network and topological statistics of the of Hm_net in comparison to single species networks. Topological statistics (*) are calculated for the largest component (LC) only.

|  | *H. sapiens* | *S. cerevisiae* | *C. elegans* | Hm_net |
| --- | --- | --- | --- | --- |
| Protein (all) | 17001 | 5883 | 3194 | 3474 |
| Interactions (all) | 276002 | 84277 | 5572 | 20684 |
| Connected components | 11 | 1 | 96 | 26 |
| Proteins (LC) | 16980 | 5883 | 2969 | 3408 |
| Interactions (LC) | 275991 | 84277 | 5442 | 20640 |
| Proteome coverage (%)* | 56.5 | 89.5 | 14.7 | 33.6 |
| Largest hub* | TRIM25 | NAB2 | gei-4 | Cullin 3 |
| Clustering coefficient* | 0.108 | 0.275 | 0.036 | 0.126 |
| Diameter* | 9 | 6 | 13 | 10 |
| Characteristic path length* | 3.082 | 2.483 | 4.802 | 3.837 |
| Mean number of interactions* | 32.508 | 28.651 | 3.666 | 12.113 |

The protein with the largest number of interactions in Hm_net was Cullin3 (n = 257), a protein involved in ubiquitination that has source interactions from ten eukaryotic species. The proteins with the largest number of interactions in the other species were the immune response E3 ubiquitin ligase TRIM25 in *H. sapiens* (n = 2384), the NAB2 mRNA binding protein in *S. cerevisiae* (n= 2580) and gei-4, a signal transduction protein, in *C. elegans* (n = 181). TRIM25 and NAB2 both have a large number of BioGRID interactions, 2,593 and 2,689, respectively, very few of which were lost during the scoring, mapping and integration process. By contrast gei-4 has just 181 interactions, all of which are present in the final network.

The distribution of topological parameters in all four networks were similar, with the scale of the scores reflecting the size and density of the networks (Figures 3 and S3-5). The degree (i.e. number of protein interaction) distribution of Hm_net (Figure 3) indicates that A) the network exhibits scale-free behaviour [Barabsi and Albert, 1999] in that it has a small number of highly connected proteins with the distribution obeying the power law, which is a hallmark of protein-protein interaction networks [Jeong et al., 2001, Maslov and Sneppen, 2002]. The other three networks also have scale-free behaviour, although *S. cerevisiae* to a lesser extent (Figure 3 B-D). The distribution patterns of betweenness and closeness centrality [Freeman, 1977] (measures of a protein’s topological importance) were also similar in all four networks (Figures S3 and 4), whereas the clustering coefficient [Watts and Strogatz, 1998] (i.e. degree of connectivity in a protein’s immediate neighborhood) distribution of Hm_net was more similar to that of *H. sapiens*, reflecting the larger proportion of human data contributing to the interologs (Figure S5).

**Figure 3.**
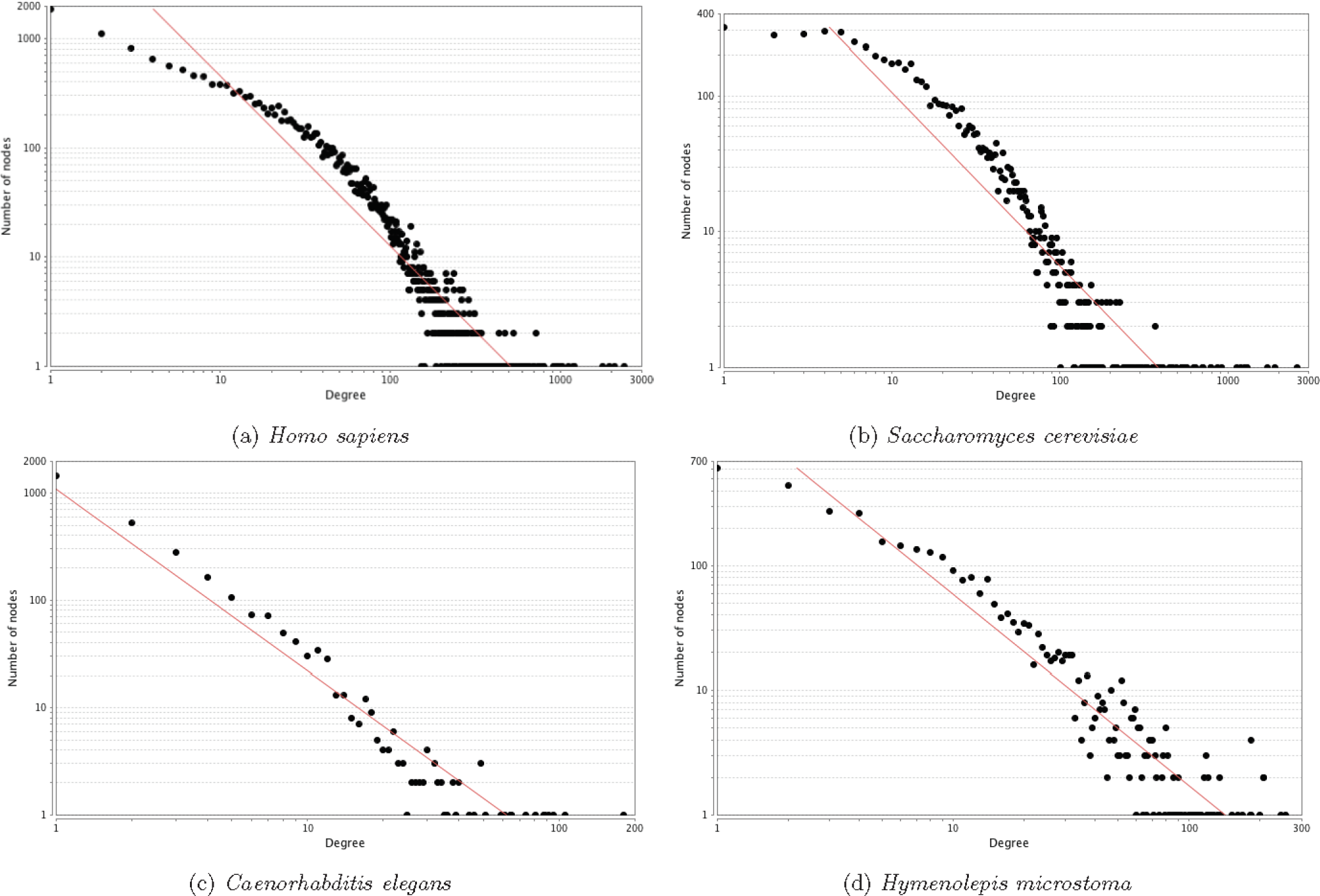
Degree distribution. The degree distribution of the four networks with the power law fitted (red). In each case the distribution is a good fit for the power law, indicating that the network has a small number of highly-interacting ‘hub’ proteins, which is a hallmark of protein-protein interaction networks: a) correlation 0.851, R-squared 0.878; b) correlation 0.660, R-squared 0.827; c) correlation 0.998, R-squared 0.902; d) correlation 0.896, R-squared 0.877.

### 3.2 Toplogically-important proteins of the high confidence network (Hm HC net) are involved in essential cellular processes

The topological statistics of a network may be used to identify the most important proteins in the network. We chose to assess the topologically-important proteins of the largest connected component of Hm HC net (1260 proteins and 3995 interactions) based on three topological scores produced by NetworkAnalyser:

1. Protein degree (number of interaction partners) to identify the top network ‘hubs’ (highly interacting proteins) [He and Zhang, 2006].
2. Betweenness centrality (BC) to identify proteins that lie between dense areas of the network [Freeman, 1977].
3. Closeness centrality (CC) to identify the most central proteins in the network in terms of information flow [Freeman, 1977].

Network hubs in protein interaction networks are often conserved and essential proteins [He and Zhang, 2006, Jeong et al., 2001, Brown and Jurisica, 2007]. The largest network hub with 88 interactions, HmN 000772200, is a cell cycle division 5-like (CDC5L) protein that is involved in the G2/M transition and known to be required for pre-mRNA splicing. The second largest hub, HmN 000015300, was also a putitive pre-mRNA processing factor. Of the remaining top network hubs, a large number of the proteins were involved in gene expression; eight ribosomal proteins; two translation initiation factors; one RNA polymerase subunit, and a histone deacetylase (Table 2). Five of the hubs were cullin family protein which play an intrinsic role in post-translational modification of protein via ubiquitination

**Table 2.** Network hubs. The top twenty-five network hubs ranked by degree (number of interactions). BC: betweenness centrality; CC: closeness centrality.

| Gene model | Degree | BC | CC | Description |
| --- | --- | --- | --- | --- |
| HmN_000772200 | 88 | 0.131 | 0.284 | Cell division cycle 5 protein |
| HmN_000015300 | 79 | 0.101 | 0.283 | Pre-mRNA processing factor 19 |
| HmN_000799800 | 52 | 0.129 | 0.297 | Heat shock protein 90 alpha |
| HmN_003006230 | 46 | 0.135 | 0.290 | DNA-directed RNA polymerase subunit RBP1 |
| HmN_000008200 | 42 | 0.046 | 0.262 | Cullin 3 |
| HmN_000077700 | 41 | 0.007 | 0.231 | 40S ribosomal protein S3a |
| HmN_000217500 | 41 | 0.004 | 0.234 | 40S ribosomal protein S2 |
| HmN_002028700 | 39 | 0.006 | 0.247 | Probable U2 small nuclear ribonucleoprotein A' |
| HmN_003003820 | 39 | 0.034 | 0.268 | Eukaryotic initiation factor 4A-III |
| HmN_003003830 | 39 | 0.034 | 0.268 | Eukaryotic initiation factor 4A-III |
| HmN_000704800 | 38 | 0.135 | 0.315 | Cyclin-dependent kinase 2 |
| HmN_000755100 | 38 | 0.019 | 0.245 | Microfibrillar associated protein 1 |
| HmN_000632800 | 38 | 0.004 | 0.243 | 116 kDa U5 small nuclear ribonucleoprotein component |
| HmN_000018400 | 37 | 0.069 | 0.255 | Histone deacetylase 1 |
| HmN_000932000 | 37 | 0.006 | 0.232 | 40S ribosomal protein S16 |
| HmN_000002900 | 37 | 0.001 | 0.223 | 40S ribosomal protein S4 |
| HmN_003026960 | 36 | 0.002 | 0.231 | 40S ribosomal protein S8 |
| HmN_000899300 | 35 | 0.001 | 0.222 | 40S ribosomal protein S13 |
| HmN_000632900 | 35 | 0.001 | 0.222 | 40S ribosomal protein S23 |
| HmN_000063500 | 34 | 0.014 | 0.269 | Cullin 1 |
| HmN_000629900 | 34 | 0.014 | 0.269 | Cullin 1 |
| HmN_003003610 | 34 | 0.014 | 0.269 | Cullin 1 |
| HmN_003003620 | 34 | 0.014 | 0.269 | Cullin 1 |
| HmN_000144900 | 33 | 0.068 | 0.302 | Cyclin-dependent kinase 2 |
| HmN_000490400 | 33 | 0.001 | 0.231 | 60S ribosomal protein L4 |

Centrality statistics measure a protein’s importance to information flow through the network [Freeman, 1977]. Betweenness centrality is a measure of the amount of influence a protein has on information flow based on the number of shortest paths between protein pairs on which it lies. High scoring proteins that also have low degree (few interactions), termed ‘bottlenecks’, are often highly conserved and essential [Joy et al., 2005, Yu et al., 2007]. Closeness centrality also measures information flow through a protein based on how short the shortest paths are from that protein to all other proteins in the network. A high score indicates the ability to communicate with other network members through a small number of intermediaries and has been used to identify key components of metabolic pathways [Ma and Zeng, 2003].

Of the top scoring proteins for betweenness centrality (Table 3) ten are also network hubs. The other fifteen proteins are six involved in the cell cycle and replication, three cytoskeletal, two histone-related, two ubiquitination, one clarthrin chain and one proteasomal protein. Four of the cell cycle proteins are transitional endoplasmic reticulum ATPases (HmN 000846600, HmN 003022520 and HmN 003022580) have interactions resulting from the same interolog evidence and, therefore, identical interactions and topological statistics in the network. The majority of high closeness centrality proteins (20 of 25) are hubs, high betweenness or both (Table 4). The remaining high CC are a splicing factor, a chaperonin, a SNW domain containing transcriptional protein, and two proteins involved in the cell cycle and replication.

**Table 3.** Betweenness centrality. The top twenty-five protein ranked by betweenness centrality. Proteins that are also top 20 protein hubs (Table 2) are denoted with *. BC: betweenness centrality; CC: closeness centrality.

| Gene model | Degree | BC | CC | Description |
| --- | --- | --- | --- | --- |
| HmN_003006230 | 46* | 0.135 | 0.290 | DNA-directed RNA polymerase subunit RBP1 |
| HmN_000704800 | 38* | 0.135 | 0.315 | Cyclin-dependent kinase 2 |
| HmN_000772200 | 88* | 0.131 | 0.284 | Cell division cycle 5 protein |
| HmN_000799800 | 52* | 0.129 | 0.297 | Heat shock protein 71 kDa protein 90 alpha |
| HmN_000015300 | 79* | 0.101 | 0.283 | Pre-mRNA processing factor 19 |
| HmN_003048860 | 24 | 0.097 | 0.277 | Actin |
| HmN_003006810 | 28 | 0.074 | 0.281 | Histone acetyltransferase p300 |
| HmN_000018400 | 37* | 0.069 | 0.255 | Histone deacetylase 1 |
| HmN_000144900 | 33* | 0.068 | 0.302 | Cyclin-dependent kinase 2 |
| HmN_002231000 | 32 | 0.051 | 0.260 | 26S proteasome non-ATPase regulatory subunit 2 |
| HmN_000536300 | 24 | 0.047 | 0.265 | Tubulin gamma chain-1 |
| HmN_000547600 | 12 | 0.047 | 0.267 | Histone H2B |
| HmN_000008200 | 42* | 0.046 | 0.262 | Cullin 3 |
| HmN_000237800 | 21 | 0.043 | 0.254 | Dual specificity protein phosphatase cdc14a |
| HmN_000405000 | 14 | 0.038 | 0.258 | Clathrin heavy chain 1 |
| HmN_000846600 | 30 | 0.037 | 0.288 | Transitional endoplasmic reticulum ATPase |
| HmN_003022520 | 30 | 0.037 | 0.288 | Transitional endoplasmic reticulum ATPase |
| HmN_003022580 | 30 | 0.037 | 0.288 | Transitional endoplasmic reticulum ATPase |
| HmN_002012300 | 10 | 0.036 | 0.241 | Ubiquitin-conjugating enzyme E2 variant 3 |
| HmN_003003820 | 39* | 0.034 | 0.268 | Eukaryotic initiation factor 4A-III |
| HmN_003003830 | 39* | 0.034 | 0.268 | Eukaryotic initiation factor 4A-III |
| HmN_000066300 | 23 | 0.033 | 0.259 | E3 ubiquitin ligase RING3 |
| HmN_003005250 | 9 | 0.033 | 0.234 | Dynein heavy chain |
| HmN_000150200 | 23 | 0.031 | 0.256 | Proliferating cell nuclear antigen |
| HmN_000372000 | 17 | 0.030 | 0.271 | DNA replication licensing factor MCM2 |

**Table 4.** Closeness centrality. The top twenty-five protein ranked by closeness centrality. Proteins that are also top 20 protein hubs (Table 2) are denoted with * and top 20 betweenness centrality (Table 3) with **. BC: betweenness centrality; CC: closeness centrality.

| Gene model | Degree | BC | CC | Description |
| --- | --- | --- | --- | --- |
| HmN_000704800 | 38* | 0.135** | 0.315 | Cyclin-dependent kinase 2 |
| HmN_000144900 | 33* | 0.068** | 0.302 | Cyclin-dependent kinase 2 |
| HmN_000799800 | 52* | 0.129** | 0.297 | Heat shock protein 71 kDa protein 90 alpha |
| HmN_003006230 | 46* | 0.135** | 0.290 | DNA-directed RNA polymerase subunit RBP1 |
| HmN_000846600 | 30 | 0.037** | 0.288 | Transitional endoplasmic reticulum ATPase |
| HmN_003022520 | 30 | 0.037** | 0.288 | Transitional endoplasmic reticulum ATPase |
| HmN_003022580 | 30 | 0.037** | 0.288 | Transitional endoplasmic reticulum ATPase |
| HmN_000772200 | 88* | 0.131** | 0.284 | Cell division cycle 5 protein |
| HmN_000015300 | 79* | 0.101** | 0.283 | Pre mRNA processing factor 19 |
| HmN_003006810 | 28 | 0.074** | 0.281 | Histone acetyltransferase p300 |
| HmN_003048860 | 24 | 0.097** | 0.277 | Actin |
| HmN_000372000 | 17 | 0.030** | 0.271 | DNA replication licensing factor MCM2 |
| HmN_000063500 | 34* | 0.014 | 0.269 | Cullin 1 |
| HmN_000629900 | 34* | 0.014 | 0.269 | Cullin 1 |
| HmN_003003610 | 34* | 0.014 | 0.269 | Cullin 1 |
| HmN_003003620 | 34* | 0.014 | 0.269 | Cullin 1 |
| HmN_003003820 | 39* | 0.034** | 0.268 | Eukaryotic initiation factor 4A-III |
| HmN_003003830 | 39* | 0.034** | 0.268 | Eukaryotic initiation factor 4A-III |
| HmN_000547600 | 12 | 0.047** | 0.267 | Tubulin gamma chain-1 |
| HmN_000536300 | 24 | 0.047** | 0.265 | Histone H2B |
| HmN_000476900 | 23 | 0.025 | 0.265 | Splicing factor 3b subunit 3 |
| HmN_000109300 | 14 | 0.011 | 0.264 | Cyclin dependent kinase 7 |
| HmN_000744000 | 16 | 0.025 | 0.263 | Chaperonin containing TCP1 subunit 2 (beta) |
| HmN_000541900 | 13 | 0.018 | 0.262 | Replication protein A 70 kDa DNA-binding subunit |
| HmN_000463400 | 22 | 0.028 | 0.262 | SNW domain containing protein 1 |

### 3.3 Network clusters of the high confidence network (Hm HC net) correspond to biological modules and processes

We used the MCODE algorithm to identify tightly connected area of the the largest component of Hm HC net since clusters in protein-protein networks from model species generally correspond to complexes of proteins involved in the same biological process [Girvan and Newman, 2002]. A total of 38 clusters were identified in Hm_net, ranging from MCODE score 26.5 to 2.7, and 3 to 27 proteins in size.

The ten highest scoring clusters represent proteins with related biological functions (Figures 7 and S6; Tables 5). The largest and highest scoring cluster comprises 27 proteins, 26 of which are ribosomal subunits in addition to a single proteasome subunit, HmN 000306800. Clusters 2 and 3 represent groups of proteaosme and DNA-directed RNAP proteins, respectively, with the exception of HmN 003000770 in cluster 3, which is a SWI/SNF-related chromatin regulator. Cluster 4 contains 11 proteins related to mRNA processing and the spliceosome. Cluster 5 contains proteins that are mainly related to microtubules, 8 tubulins, dynactin and dystonin, in addition to an ADP-ribosylation factor and two unannotated proteins, HmN 000742700 and HmN 000742800.

**Table 5.** Network clusters. Cluster metrics for the ten clusters shown in Figure 7. Cluster descriptions are based on the majority of annotations to the cluster proteins; full cluster annotations are provided in Table S4.

| Cluster | MCODE Score | Proteins | Interactions | Description |
| --- | --- | --- | --- | --- |
| 1 | 26.462 | 27 | 344 | Ribosome |
| 2 | 11.667 | 13 | 70 | DNA-dependent RNA polymerase |
| 3 | 7.429 | 8 | 26 | Proeasome |
| 4 | 7.200 | 11 | 36 | Spliceosome |
| 5 | 6.667 | 13 | 40 | Tubulins |
| 6 | 5.250 | 9 | 21 | Translation initiation complex |
| 7 | 5.000 | 5 | 10 | snRNP binding (LSm) |
| 8 | 5.000 | 9 | 20 | 14-3-3 proteins |
| 9 | 5.000 | 9 | 20 | PAK kinases |
| 10 | 5.000 | 9 | 20 | RhoA |

**Figure 4.**
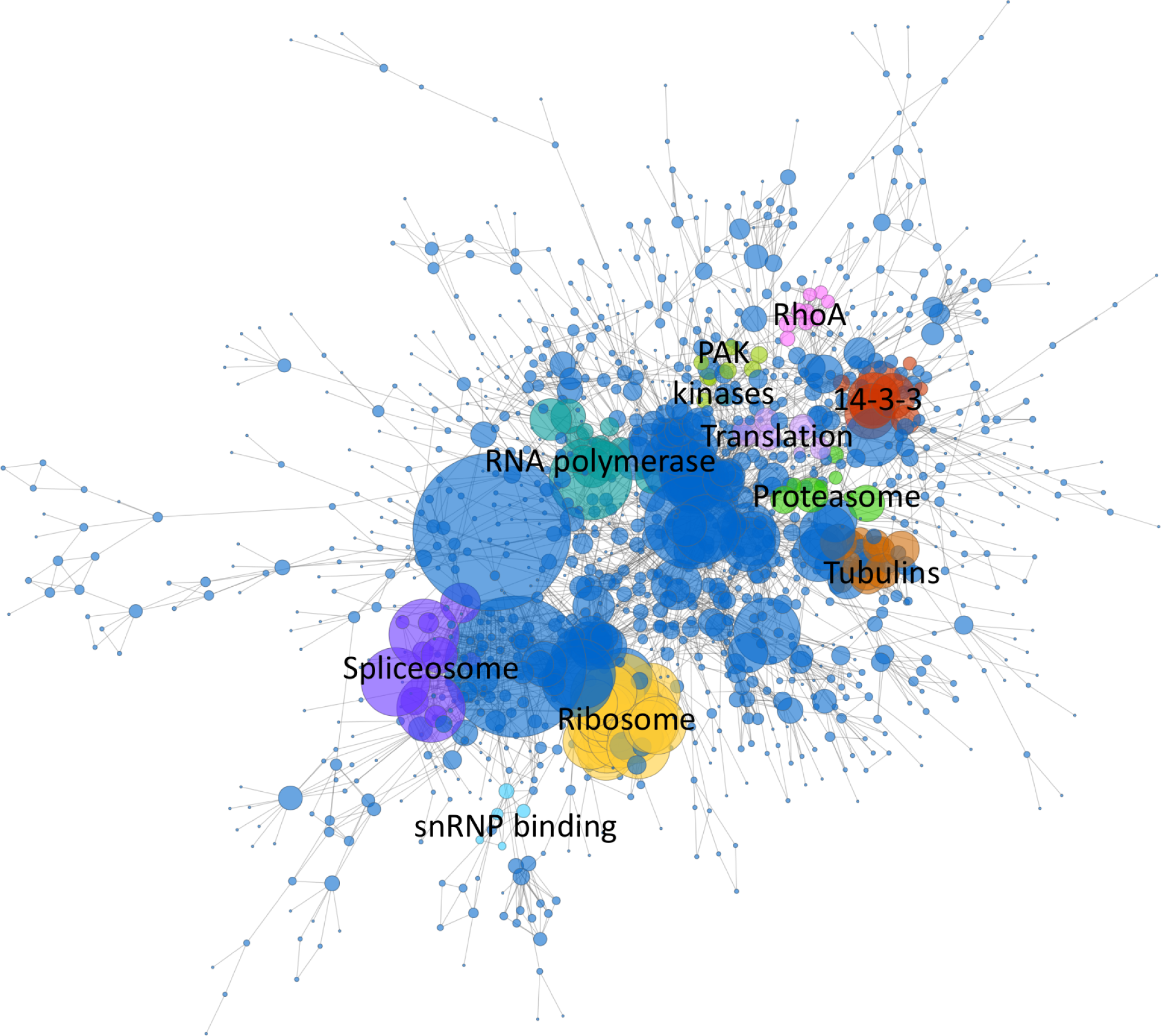
Network clusters. The ten highest scoring MCODE clusters are highlighted in Hm_net. Protein nodes are sized by number of interactions. Cluster 1: yellow, cluster 2: turquoise, cluster 3: green, cluster 4: purple, cluster 5: brown, cluster 6: mauve, cluster 7: light blue, cluster 8: red, cluster 9: pale green, cluster 10: pink. Full cluster annotations are provided in Table S4.

**Figure 5.**
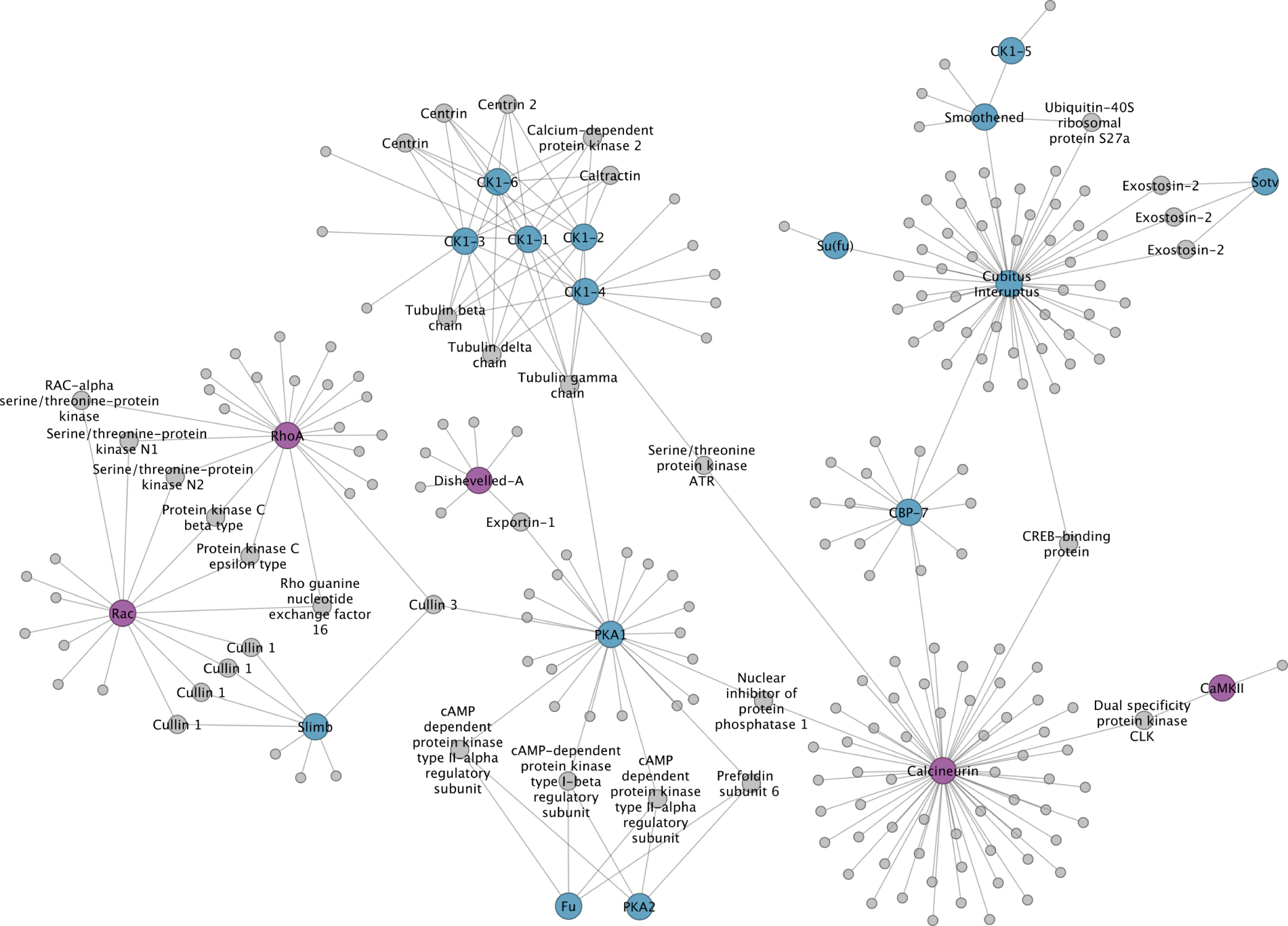
Hedgehog and Wnt intermediaries.

**Figure 6.**
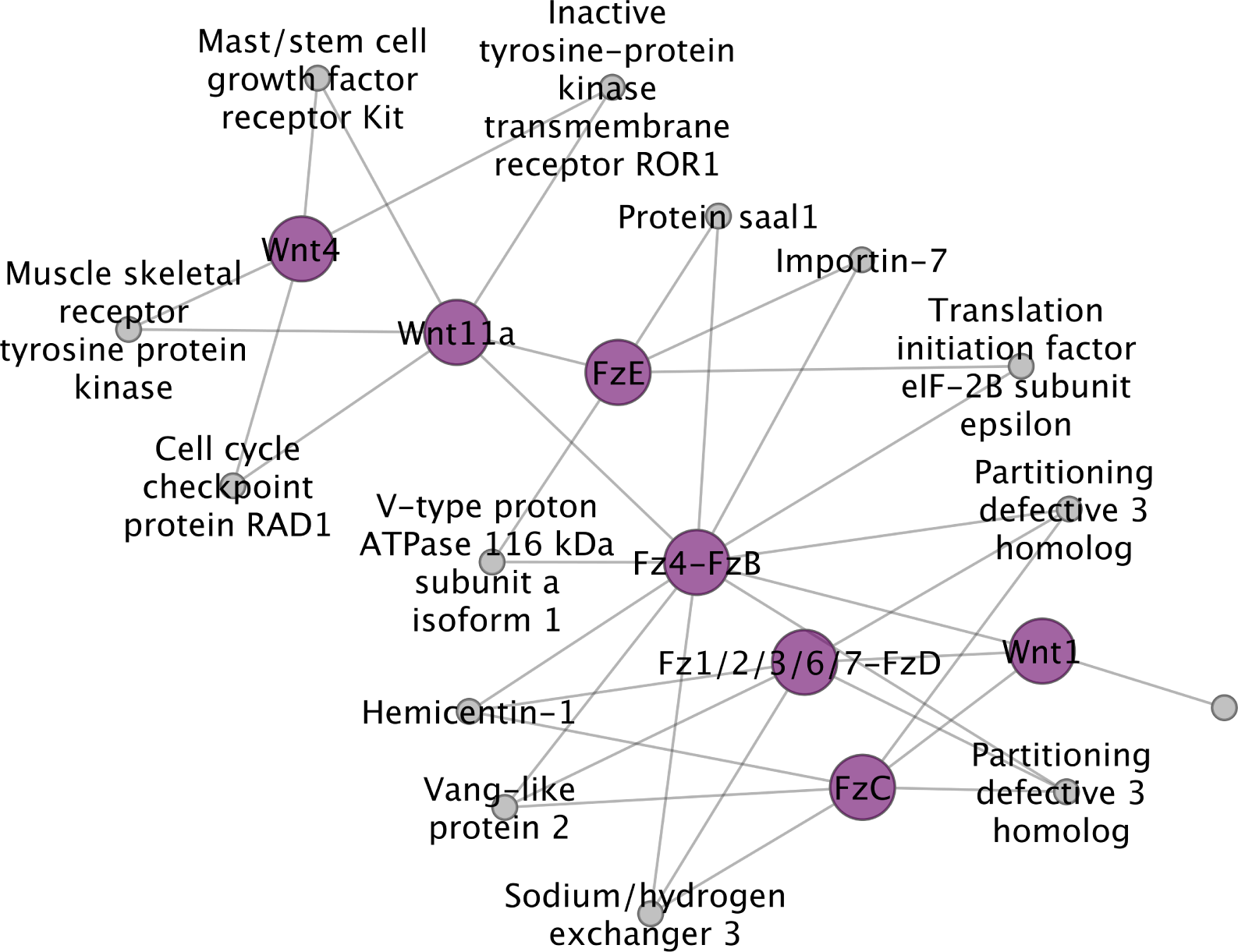
Wnt ligands and Frizzled receptors.

**Figure 7.**
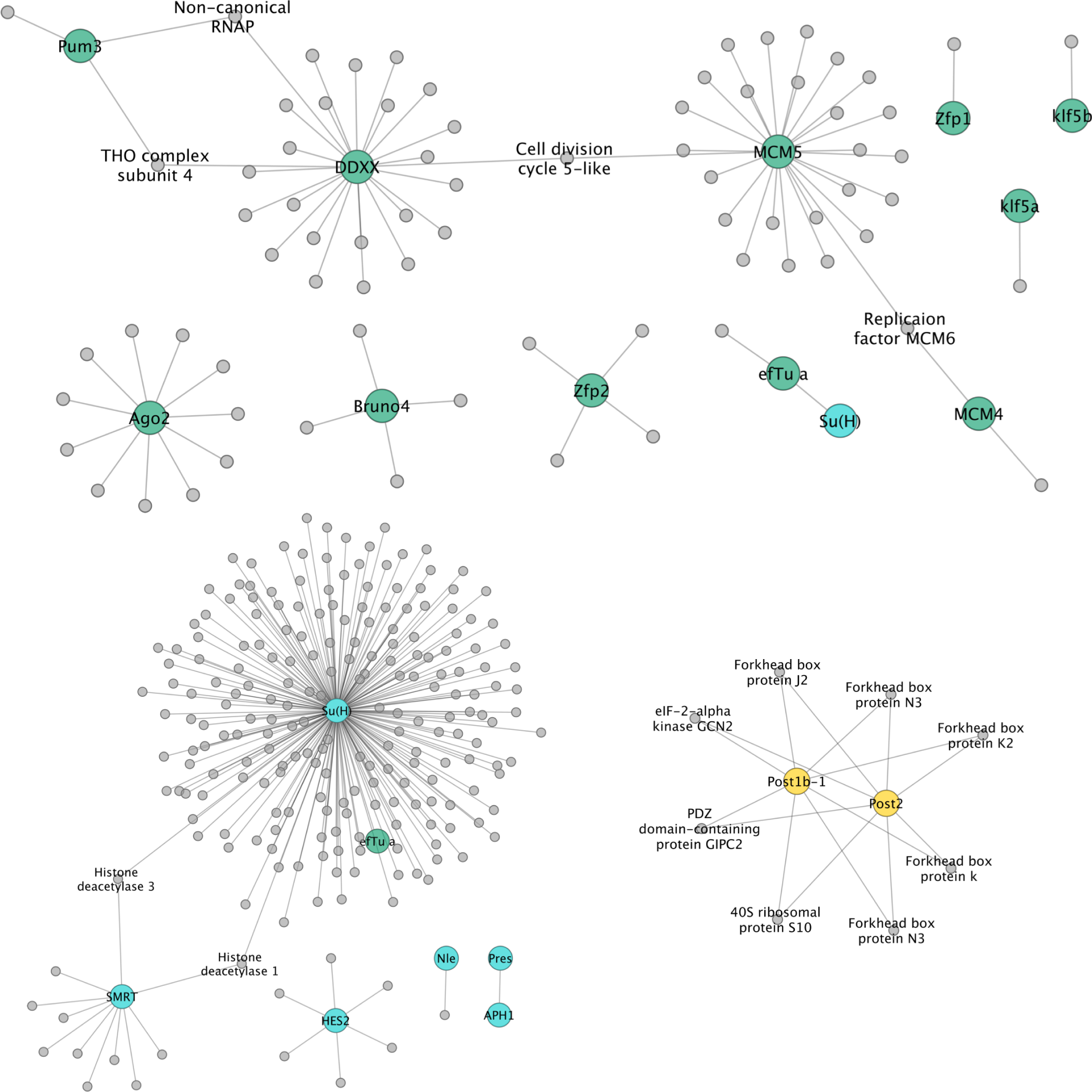
Germline ‘multipotency’ genes, Notch intermediaries and Hox genes.

Cluster 6 comprises six eukaryotic translation initiation factors and two subunits of the COP9 signalosome, a regulator of the Cullin-RING proteins. Cluster 7 contains four of the Lsm proteins which bind U6 snRNPs and a probable snRNP. Finally, clusters 8, 9 and 10 are signalling and regulatory clusters each comprising several kinases in addition to members of the 14-3-3 proteins, PAK kinases and RhoA pathways, respectively. Full cluster annotations are provided in Table S4.

### 3.4 Specific genes of interest: signalling, transcription and germline-related genes

Probably the most common use of interactome prediction is to identify proteins that, based on knowledge in other organisms, are likely to be associated with specific genes of interest (GOI). To this end, we examined specific suites of proteins representing components of select signalling pathways, transcription factors and germline/stem-cell-related multipotency genes. The results of these are shown in Figures 5-7. As expected, the majority of the GOIs that were present in the network, and that also showed the largest number of interactions, were signal transduction components or cell cycle regulators that are highly conserved and among the former, often operate across multiple pathways. This is illustrated in Figure 5 in which intracellular components of Wnt and Hedgehog signalling are predicted to be connected functionally by way of cullin proteins (which help direct ubiquitin-mediated protein destruction), protein kinases (involved in phosphorylation), and additional factors. Among these are some of the most highly connected proteins among the GOIs, such as RhoA, a hydrolase that acts in the Wnt planar cell polarity pathway and Calcineurin, a protein phosphatase involved in dephosphorylation which acts in the calcium-dependent Wnt pathway (Figure 5). Additional examples are Cubitus interruptus, a zinc finger transcription factor responsible for activating downstream target genes (such as Wnt1) in Hedgehog signalling, and Suppressor of hairy, a bi-functional protein that mediates activation or repression of other proteins in the Notch signalling pathway (Figure 7).

Figure 6 shows interactions between and within Wnt ligands and their canonical receptors, the frizzled transmembrane proteins. Results predict direct interactions between the posterior morphogen Wnt1 and three of the five frizzled receptors in their genome (fz4, fz1/2/3/6/7 and fzC) [Riddiford and Olson, 2011]. The posterior Wnt11 ortholog Wnt11a is also predicted to interact with fz4, but also with fzE (see S3 for corresponding gene models). Wnt4, by contrast, is not linked to a frizzled protein. Among interactions with other proteins, links between Wnt4 and Wnt11a to the cell cycle checkpoint protein Rad1 may be one means by which the canonical, *β*-catenin-dependent, Wnt pathway can regulate cell proliferation and thus growth.

Figure 7 shows germline, or stem-cell, multipotency proteins, components of the Notch signalling pathway (discussed above) and Hox transcription factors found in Hm_net. Relatively few putative stem-cell related proteins were found in the network, but those that were are connected, as expected, by other regulators of the cell cycle such as CDC5-like. Similarly, most predicted interactions are with housekeeping or cell cycle regulatory genes. However, one of two zinc finger transcription factors putatively associated with flatworm stem cells [van Wolfswinkel et al., 2014] has four predicted interacting proteins, all of which are SMAD factors, the intracellular transducers of the TGF*β*/BMP signalling pathways.

Only two Hox family transcription factors were found to be present in the network: a bona fide Post2/Abda ortholog and a Post1/Post2-like posterior Hox paralog [Olson, 2008]. Although the sequences of these two proteins are very divergent, both are annotated with identical interacting proteins, and thus would be predicted to be playing the same role in the organism. Interestingly, five of the eight associated proteins are forkhead box (FOX) transcription factors, which have been shown to interact with Hox genes [Brayer et al., 2011], and thus provides predictions of which of the numerous FOX proteins in tapeworms to investigate in relation to Hox expression.

### 3.5 Differentially expressed genes: comparing the interactomes of larval and adult worms

We were interested in exploring interactomes specific to different life stages. We mapped differentially-expressed genes (DEGs) identified between adults and 5-day old, metamorphosing larvae [Olson et al., 2018] to the network to create a sub-network of these proteins (where the DEGs are present in the network). Of 3,479 DEGs, 367 were present in the network, and 176 proteins formed a connected component of 668 interactions (Figure 8). Several members of the network clusters (Figure 7) were represented in the DEG subnetwork.

**Figure 8.**
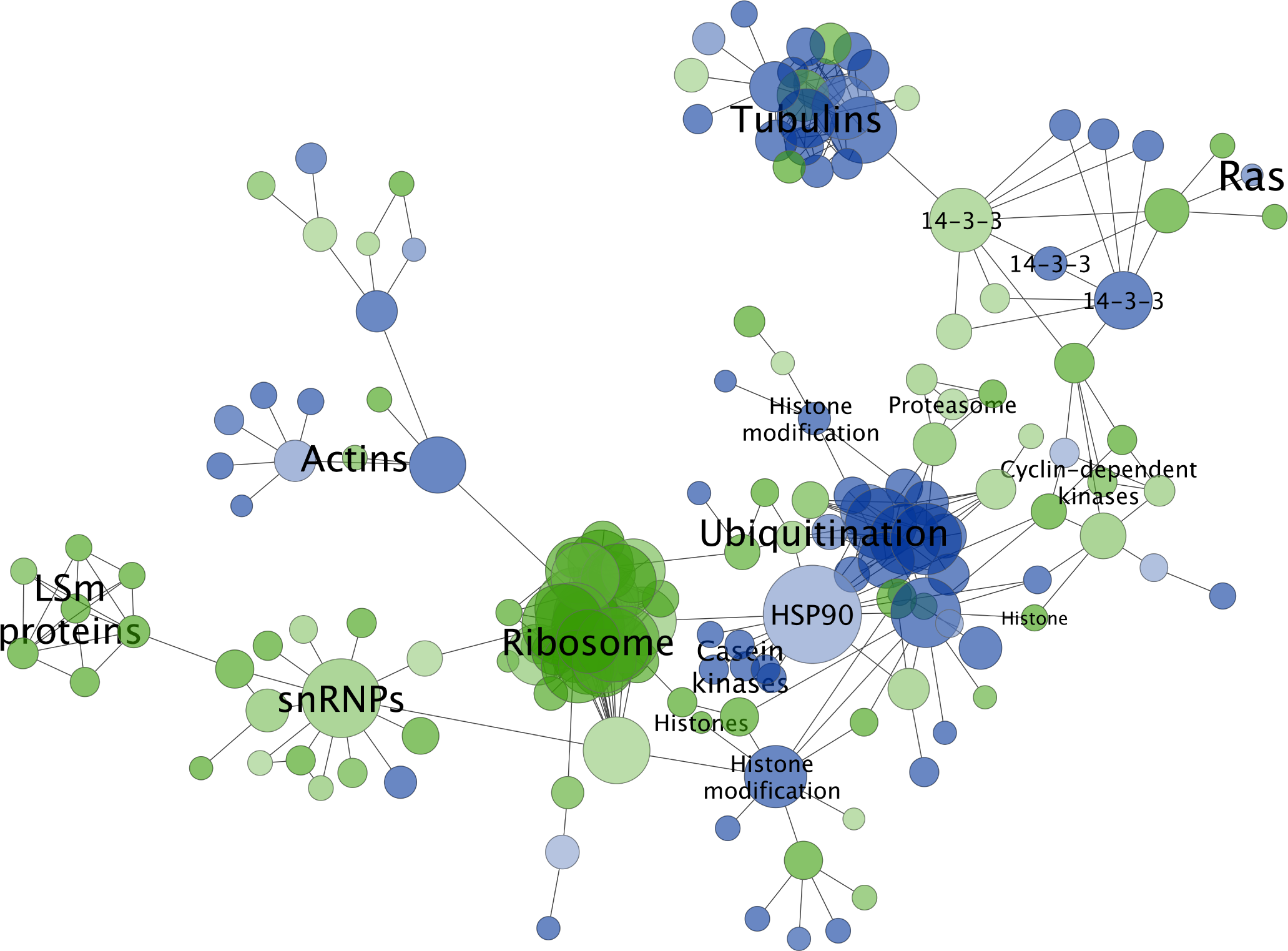
Differentially-expressed genes between adult and larval worms were mapped to Hm_HC_net and the sub-network extracted (green, up-regulated in larvae; blue, up-regulated in adults). Protein nodes are sized relative to the number of interactions they connect.

In larvae, ribosomal and RNA splicing factors were significantly up-regulated. These proteins included a large ribosomal group corresponding to 23 of 27 proteins of cluster 1 in addition to twelve other ribosomal proteins. This ribosomal group was connected to a group of thirteen snRNP-related proteins that contained three proteins of cluster 4, which was, in turn, connected to the Lsm proteins of cluster 7. Four of the eight proteasomal subunits of cluster 2 were also up-regulated in larvae. A group of three histones are also up-regulated.

In adults, two groups of cytoskeletal proteins (actins and tubulins) were up-regulated, including seven members of cluster 5, one of which HmN 000742800, is un-annotated. A large connected group of ubiquitination-associated proteins were up-regulated in adult worms, with the exception of four proteins at the periphery of the group, which had higher expression in larvae. The heat shock protein HSP90, one of the largest hubs in Hm HC net with 52 interactions, was also up-regulated in addition to a group of 6 casein kinases, and two histone modification proteins.

Signalling proteins were split between up-regulation in larvae and up-regulation in adults. The majority of cyclin-dependent kinases present in the DEG network were up-regulated in larvae, while Ras is up-regulated in adult worms. Two 14-3-3 signalling proteins were up-regulated in adults and one in larvae. Interestingly, the three 14-3-3 proteins share interaction interactions partners with one another. Full differential expression results are provided in Supplementary Table S5.

## 4 Discussion

Although a near complete chromosome-level assembly and set of stable gene models are available for *H. microstoma*, no systems-level data are available for the species. Hm_net provides the first steps toward using such data to understand tapeworm cellular biology by integrating interologs from sixteen eukaryotic species into an inferred interactome for *H. microstoma*. The network is probabilistic which reduces the impact of data set noise, in particular, from false positive interactions produced by high throughput studies, while assigning higher probabilities to interactions with multiple lines of experimental evidence [Lee et al., 2004]. Since the interologs are produced from a number of different species, these networks can have an additional level of noise, as not all interactions occurring in the source organisms will occur in *H. microstoma*. However, the confidence scoring schema also mitigates the effect of this noise in comparison to an unweighted interolog network [Lee et al., 2008]. We included a minimum count of 1.0 during the scoring stage in order to reduce loss of data in species with sparse interaction and gold standard data. While this would not be necessary or desirable in well studied species, the thresholding of the network to remove low scoring interactions allows for the retention of those with multiple lines of low scoring evidence, which would otherwise have been lost without the minimum count.

The topologically-important nodes and clusters of Hm_net represent core housekeeping and essential processes, which is to be expected as these processes are common to all the species from which the interlogs are derived. For example, the COP9 signalosome is found in all eukaryotes [Schwechheimer, 2004] and subunits of this complex cluster together in Hm_net. Notably, patterns of differential expression correspond to the network clustering and are connected in areas of up- and down -regulation within the network. Therefore, the network connections provide a biologically-relevant picture of *H. Microstoma* cellular biology.

It has been observed that interaction data, and annotations of proteins themselves, are biased towards certain biological processes such as protein biosynthesis and ribosomal proteins [Lee and Marcotte, 2009, Myers et al., 2006], so it is unsurprising that Hm_net shows similar biases. The majority of significant proteins, clusters and differentially-expressed genes belong to essential, conserved processes, with the ribosomal proteins being prominent in all our assessment results. Some previous network-based studies have chosen to identify and remove these biases either during or following integration [Lee et al., 2007, Chen et al., 2005, 2006], and this approach may be of benefit when a less specific process is of interest. However, these approaches can come at the cost of the removal of valid and useful data [James et al., 2009].

One drawback of an interolog-base integration schema is the effect of redundant interologs which are based on the same evidence. *Hymenolepis microstoma* gene models with the same blast hits naturally have identical interactions and confidence scores in Hm_net. These interactions are likely to affect the results of some topological analyses, for instance, by artificially unweighting the degree of some proteins and producing tighter clustering co-efficients. In several cases these interactions were apparent during clustering and subnetwork analysis, in particular the four Cullin 1 protein hubs (Table 2) and several clusters containing symmetrical redundant interactions, such as the posterior Hox proteins. Redundancy is also likely to affect the identification of bottleneck proteins (that is, high betweeness centrality and low number of interactions). In some cases it may aid analysis by collapsing proteins and their interactions together if they have the same source evidence.

The major advantage of a network based approach is the ability to generate testable hypotheses for more focused experimental study in organisms lacking experimental data. It is noteworthy that many transcription factors are present in our networks, providing the potential to predict regulators and/or targets of genes of interest, which can be difficult from sequences alone. In addition, of particular interest are the 14-3-3 proteins that feature prominently in Hm_net as a cluster of nine proteins, three of which are found in the differential expression subnetwork (two up-regulated in adult and one in larvae) and share interaction partners. These signalling ligands are highly conserved in eukaryotes [Aitken et al., 1992] and are found in the excretory vesicles of the *Echinococcus granulosus* larvae where they are thought to modulate host immunity [Teichmann et al., 2015]. Focused study of these proteins and their shared interaction partners may aid in understanding host-parasite cross-talk [Brehm, 2010].

Another protein of interest in Hm_net is Cdc14a (HmN 000237800), which has high betweenness but a relatively low number of interactions (21). This protein is involved in cell cycle arrest and is conserved between most of the species included in the interolog network build [Saito et al., 2004, Powers and Hall, 2017, Sacristán et al., 2011]. Cdc14a may represent a ‘bottleneck protein which is likely to be essential [Joy et al., 2005, Yu et al., 2007]. Analysis of these and other network-based features has been used successfully in the prediction of essential genes across diverse organisms [Azhagesan et al., 2018]. Prediction of synthetic lethal relationships between genes is another potential network use, for instance Benstead-Hume and colleagues used protein-protein interaction networks to predict human synthetic lethal interactions, which they then confirmed experimentally [Benstead-Hume et al., 2019]. Such analyses could potentially be used to identify targets for new chemotherapies in helminth research [Pinto et al., 2014].

Networks may also be used to predict protein function based on interaction patterns, which is especially useful where there is no sequence similarity to other known proteins [Sharan et al., 2007]. For example, HmN 000742700 and HmN 000742800, although un-annotated, cluster in the network with the tubulins (Figure 7). Additionally, HmN 000742800 shares an expression profile with a large group of connected tubulin proteins (Figure 8), making it a candidate for involvement on tubulin-related processes.

A final potential use of this network is in comparative interactomics with other species in terms of presence/absense of interologs. Network comparison has the potential to identify areas of conservation and of divergence in interaction patterns [Liang et al., 2006]. The caveat to this type of approach is that the proteome of the comparison species must be as complete as that of *Hymenolepis microstoma*, otherwise differences observed may not necessarily be biological. However, both the human bloodfluke *Schistosoma mansoni* and the tapeworm *Echinococcus multilocularis* have equally complete proteomes, providing the potential for cross-species comparison.

We note that the network is far from complete in terms of proteome coverage but covers a larger proportion of the proteome than the equivalent network for the model worm *C. elegans*. In fact, the number of interactions for *C. elegans* is low, in comparison to the other model species, which is likely due to there only being two HTP datasets available [Li et al., 2004, Simonis et al., 2009]. Inclusion of other types of interaction has the potential to increase this coverage of the *Hymenolepis microstoma* proteome. For example, regulogs networks link orthologs of regulatory interactions [Yu et al., 2004] and associalog networks link proteins/genes based on any type of interaction: physical, genetic, regulatory and other types of functional association [Kim et al., 2013, Lee et al., 2008, Shim et al., 2017]. However, these approaches generally come at the cost of more noise from false positives [Kim et al., 2013, Lee et al., 2008].

Experimental demonstration of protein-protein interactions can require considerable effort and so far no high-throughput approach has been applied to parasitic flatworms. In a new study, Montagne and colleagues used a yeast two-hybrid system and additional means to investigate the downstream effectors of canonical Wnt signalling in tapeworms, showing that only one of three paralogs of *β*-catenin actually interacts with components of the canonical destruction pathway [Montagne et al., 2019], similar to the situation in free-living planarians [Su et al., 2017]. This represents one of the first such studies to test protein interactions in tapeworms, illustrating the scarcity of experimental data available for these important pathogens. With complete genomes now available, the application of systems level analyses can start to play an important role in ameliorating this deficit by consolidating knowledge derived from major model organisms. To help achieve this, in future studies we will expand Hm_net to include regulogs and associalogs, and perform comparative interactomics between *Hymenolepis microstoma* and other helminth species.

## Conflict of Interest Statement

The authors declare that the research was conducted in the absence of any commercial or financial relationships that could be construed as a potential conflict of interest.

## Author Contributions

KJ conceived and executed the study and led manuscript preparation. PDO contributed to its inception, interpretation and writing.

## Funding

The authors received no specific funding for this work.

## Acknowledgments

We thank Alan Tracey, Nancy Holroyd and Matt Berriman (Wellcome Trust Sanger Genome Institute), and Andrew Baillie (NHM), for production of the *H. microstoma* v.3 genome and annotation set.

## Supplemental Data

### Data Availability Statement

The datasets integrated for this study can be found in the BioGRID^2^ and BioSystems databases^3^.

## Footnotes

1 https://parasite.wormbase.org/Hymenolepismicrostomaprjeb124/

2 https://thebiogrid.org

3 https://www.ncbi.nlm.nih.gov/biosystems/

