## Supplemental Material for "The tapeworm interactome: inferring confidence scored protein-protein interactions from the proteome of *Hymenolepis microstoma*"

### 1 Figures

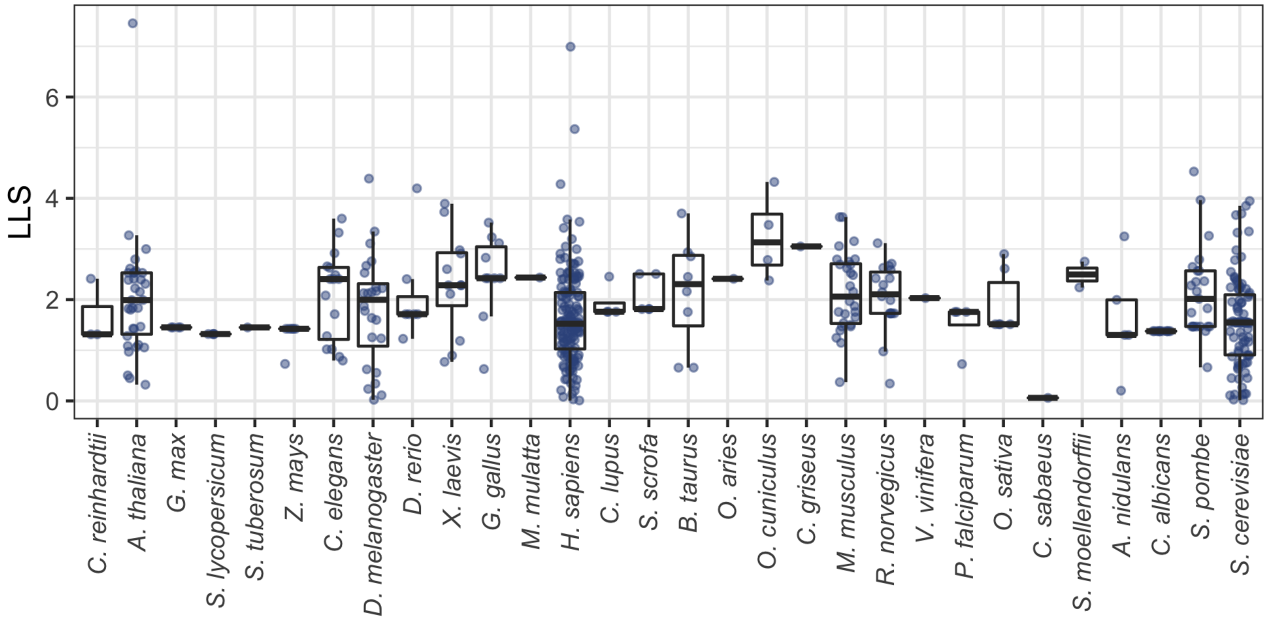


Figure S1: **Dataset confidence scoring.** The range of datasets loglikelihood (LLS) confidence scores for the BioGRID species. The majority of datasets were lost since they did not have any interactions that were interologs for *H. microstoma* gene models. Sixteen species were included in the final network (main text Table 1B).

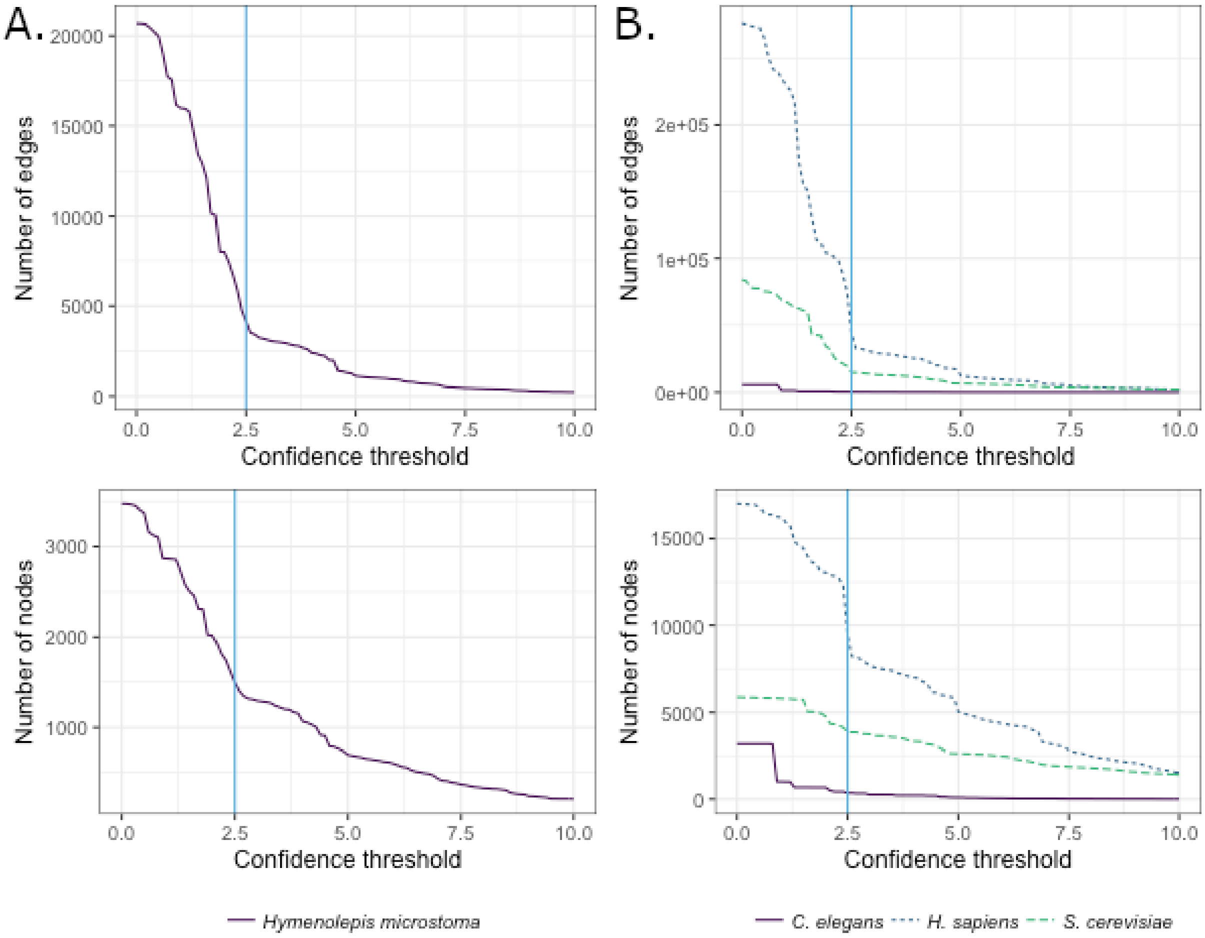


Figure S2: **Confidence thresholds.** A. Hm_net was filtered at a confidence score of 2.5 (vertical blue line), corresponding to a drop in distribution of confidence scores, to produce the high confidence Hm_HC_net. B. The *H. sapiens* and *S. cerevisiae* networks have a similar drop in confidence score distribution at a score of 2.5 despite being larger and far more densely-connected than Hm_net. The majority of *C. elegans* interaction scores are $< 1.0$.

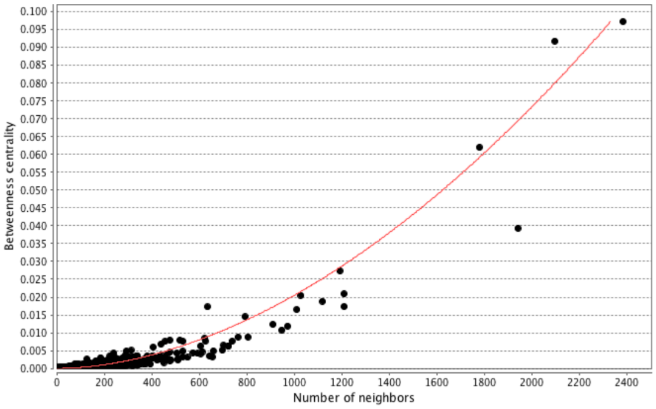


(a) *Homo sapiens*

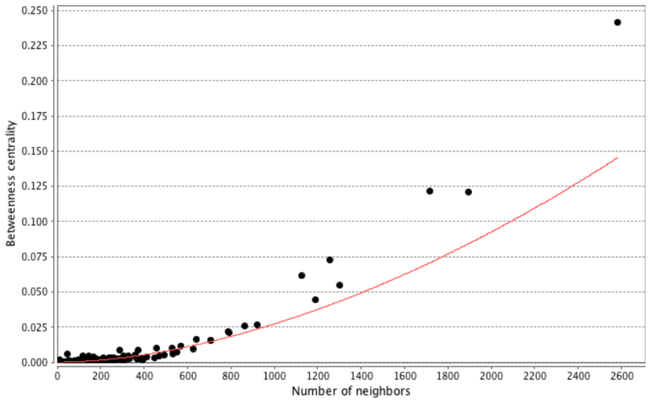


(b) *Saccharomyces cerevisiae*

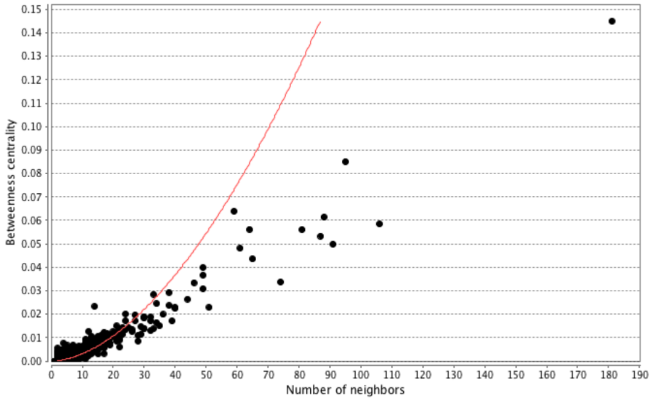


(c) *Caenorhabditis elegans*

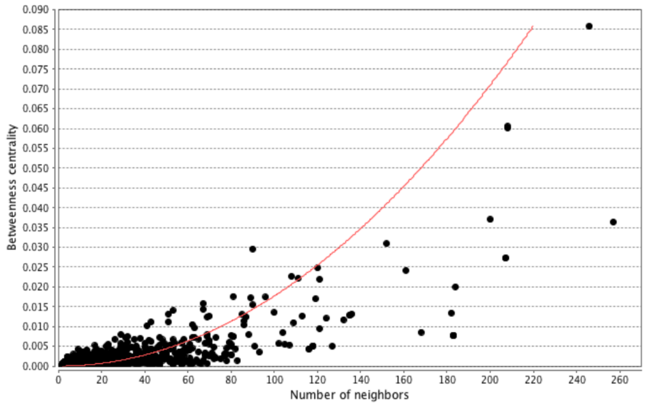


(d) *Hymenolepis microstoma*

Figure S3: **Betweenness centrality distribution.** The distribution of betweenness centrality of the four networks with the power law fitted (red). In each case the distribution is a good fit for the power law, indicating that the network has a small number of high betweenness proteins, with the majority being high betweenness.

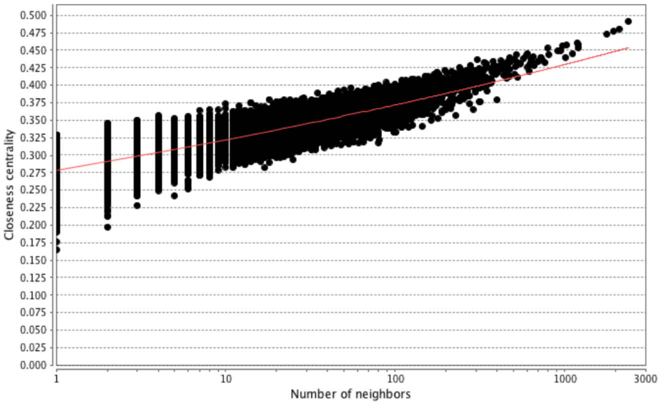


(a) *Homo sapiens*

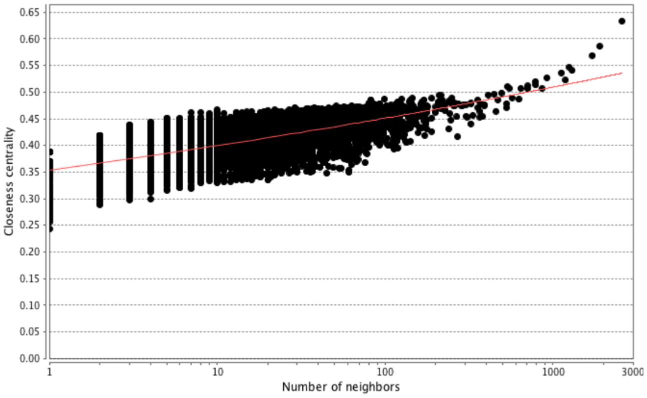


(b) *Saccharomyces cerevisiae*

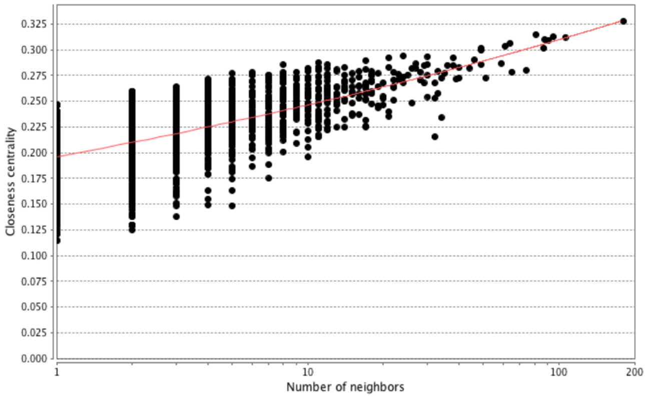


(c) *Caenorhabditis elegans*

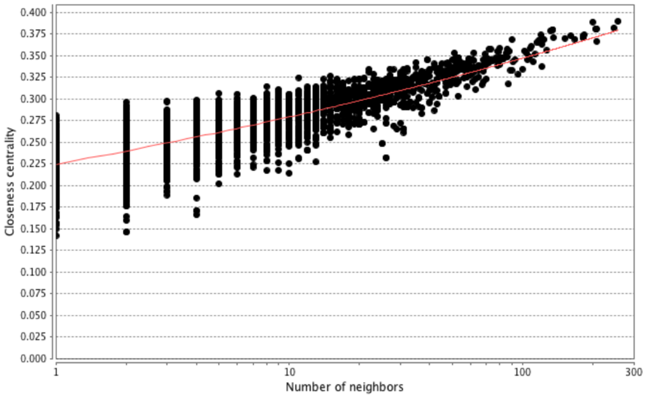


(d) *Hymenolepis microstoma*

Figure S4: **Closeness centrality distribution.** The distribution of closeness centrality of the four networks with the power law fitted (red). In each case the distribution is a good fit for the power law, indicating that the network has a small number of high closeness proteins, with the majority being high closeness.

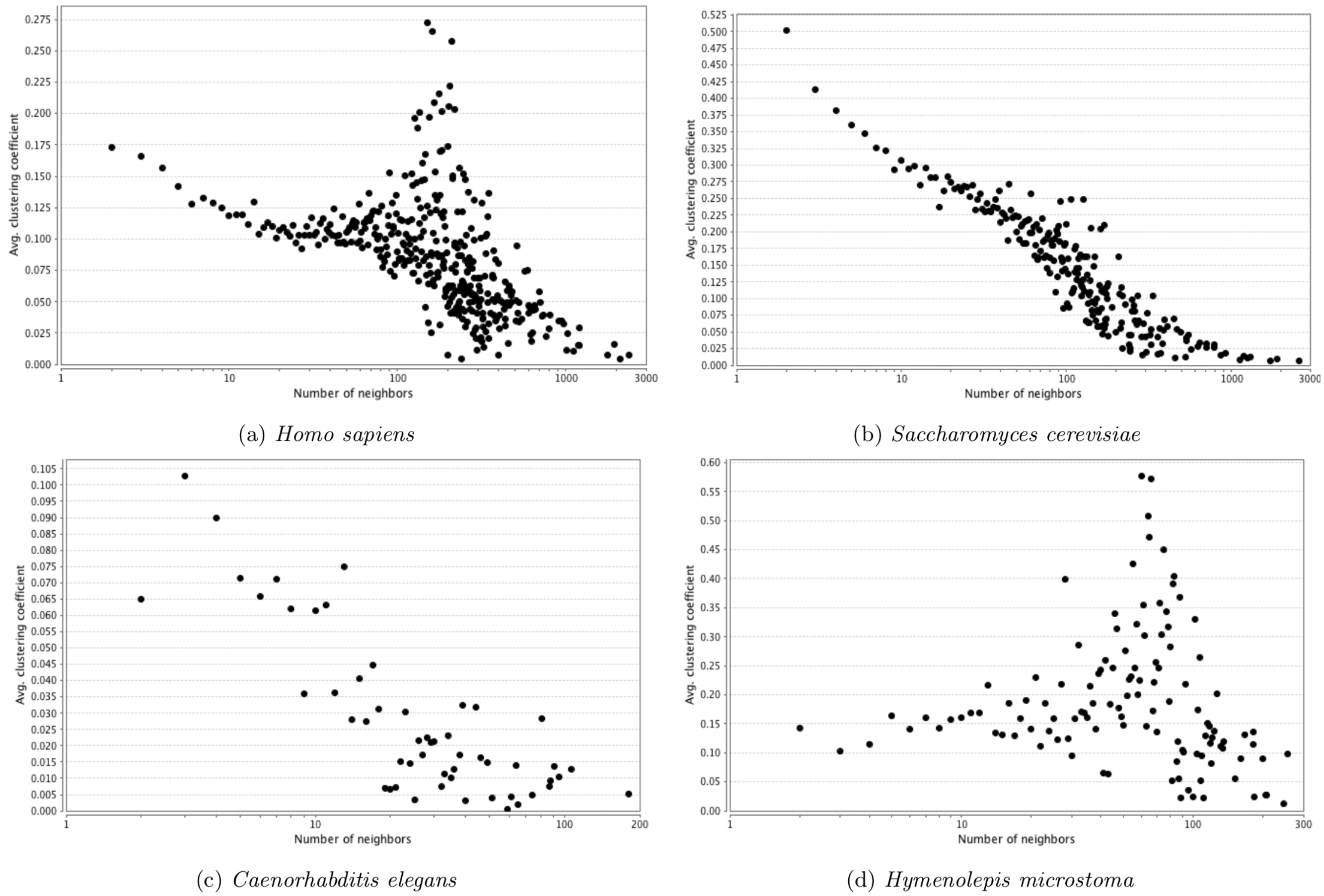


(a) *Homo sapiens*

(b) *Saccharomyces cerevisiae*

(c) *Caenorhabditis elegans*

(d) *Hymenolepis microstoma*

Figure S5: **Clustering coefficient distribution.** The distribution of clustering coefficient of the four networks. In general, the proteins with a higher number of neighbours have a higher clustering co-efficient in all four cases. The distribution of Hm_net is most similar to that of *H. sapiens*, in which a group of proteins with a lower number of neighbours form a high clustering co-efficient peak, likely reflecting the larger proportion of human data contributing to the interologs.

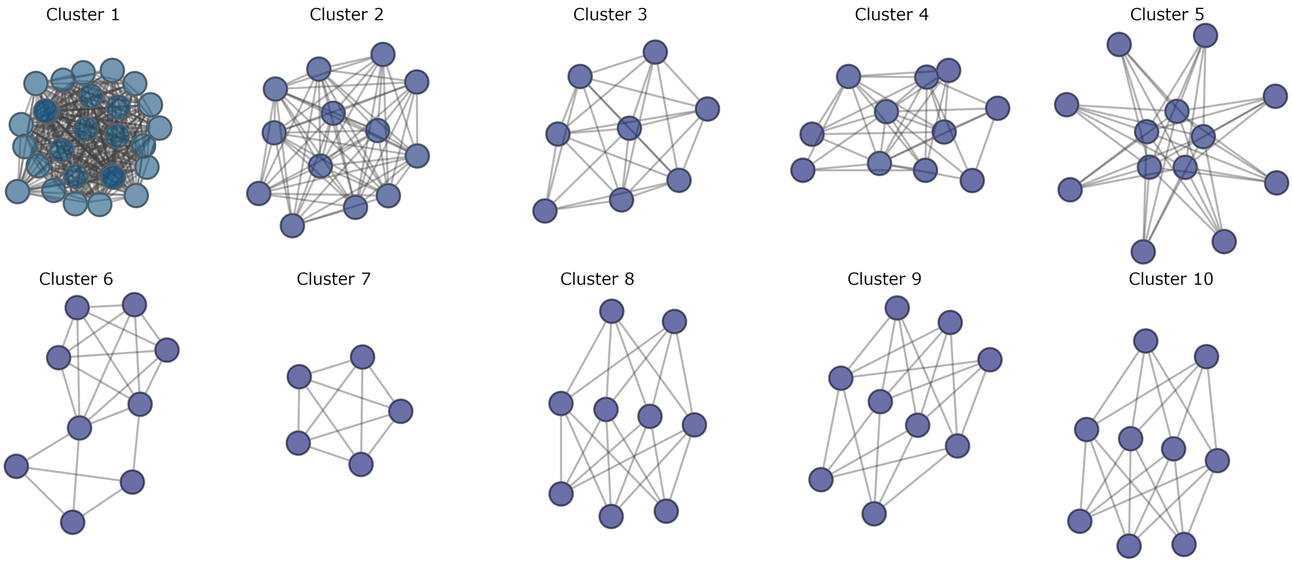


Figure S6: **Network clusters.** The ten largest network clusters (Tables S4, main text Table 5 and Figure 4): cluster 1 - ribosome; cluster 2 - DNA-dependent RNA polymerase; cluster 3 - proteasome; cluster 4 - spliceosome; cluster 5 - tubulins; cluster 6 - translation initiation complex; cluster 7 - snRNP binding; cluster 8 - 14-3-3 signaling; cluster 9 - PAK kinases; cluster 10 - RhoA.

### 2 Tables

Table S1: **Low throughput data.** The low throughput (< 200 interactions) datasets extracted from V164 of the BioGRID database. Interaction data were limited to physical interactions from eukaryotic species then split into individual datasets by study and species. Low throughput data were then grouped into datasets by BioGRID experimental type.

| **Species** | **Type** | **Interactions** |
|---|---|---|
| *Chlamydomonas reinhardtii* | Affinity.Capture-MS | 11 |
| | Biochemical Activity | 1 |
| | Reconstituted Complex | 2 |
| *Arabidopsis thaliana* | Affinity Capture-Luminescence | 46 |
| | Affinity Capture-MS | 1715 |
| | Affinity Capture-RNA | 21 |
| | Affinity Capture-Western | 1364 |
| | Biochemical Activity | 443 |
| | Co-crystal Structure | 45 |
| | Co-fractionation | 118 |
| | Co-localization | 58 |
| | Co-purification | 54 |
| | Far Western | 35 |
| | FRET | 569 |
| | PCA | 1168 |
| | Protein-peptide | 44 |
| | Protein-RNA | 8 |
| | Reconstituted Complex | 1682 |
| | Two-hybrid | 5020 |
| *Glycine max* | Affinity Capture-Western | 2 |
| | Co-fractionation | 1 |
| | FRET | 5 |
| | Two-hybrid | 21 |
| *Solanum lycopersicum* | Affinity Capture-Western | 2 |
| | FRET | 2 |
| | Reconstituted Complex | 10 |
| | Two-hybrid | 92 |
| *Solanum tuberosum* | Two-hybrid | 1 |
| *Zea mays* | Affinity Capture-Western | 2 |
| | Biochemical Activity | 2 |
| | Co-fractionation | 1 |
| | PCA | 1 |
| | Reconstituted Complex | 1 |
| | Two-hybrid | 2 |

Continued on next page

Table S1: **Low throughput data.** The low throughput (< 200 interactions) datasets extracted from V164 of the BioGRID database. Interaction data were limited to physical interactions from eukaryotic species then split into individual datasets by study and species. Low throughput data were then grouped into datasets by BioGRID experimental type.

| **Species** | **Type** | **Interactions** |
|---|---|---|
| *Caenorhabditis elegans* | Affinity Capture-MS | 92 |
| | Affinity Capture-Western | 196 |
| | Biochemical Activity | 19 |
| | Co-crystal Structure | 2 |
| | Co-fractionation | 4 |
| | Co-localization | 10 |
| | Co-purification | 3 |
| | Far Western | 13 |
| | FRET | 22 |
| | PCA | 4 |
| | Protein-peptide | 1 |
| | Protein-RNA | 3 |
| | Reconstituted Complex | 84 |
| | Two-hybrid | 458 |
| *Drosophila melanogaster* | Affinity Capture-MS | 2318 |
| | Affinity Capture-RNA | 472 |
| | Affinity Capture-Western | 4219 |
| | Biochemical Activity | 337 |
| | Co-crystal Structure | 71 |
| | Co-fractionation | 523 |
| | Co-localization | 96 |
| | Co-purification | 59 |
| | Far Western | 71 |
| | FRET | 74 |
| | PCA | 176 |
| | Protein-peptide | 6 |
| | Protein-RNA | 181 |
| | Reconstituted Complex | 381 |
| | Two-hybrid | 1081 |
| *Danio rerio* | Affinity Capture-MS | 1 |
| | Affinity Capture-Western | 51 |
| | Biochemical Activity | 4 |
| | Co-crystal Structure | 1 |
| | PCA | 1 |
| | Reconstituted Complex | 13 |
| | Two-hybrid | 81 |

Continued on next page

Table S1: **Low throughput data.** The low throughput (< 200 interactions) datasets extracted from V164 of the BioGRID database. Interaction data were limited to physical interactions from eukaryotic species then split into individual datasets by study and species. Low throughput data were then grouped into datasets by BioGRID experimental type.

| **Species** | **Type** | **Interactions** |
|---|---|---|
| *Xenopus laevis* | Affinity Capture-MS | 62 |
| | Affinity Capture-RNA | 3 |
| | Affinity Capture-Western | 207 |
| | Biochemical Activity | 41 |
| | Co-crystal Structure | 1 |
| | Co-fractionation | 23 |
| | Co-localization | 7 |
| | Co-purification | 1 |
| | PCA | 1 |
| | Reconstituted Complex | 75 |
| | Two-hybrid | 13 |
| *Gallus gallus* | Affinity Capture-MS | 243 |
| | Affinity Capture-Western | 69 |
| | Biochemical Activity | 3 |
| | Co-crystal Structure | 1 |
| | Co-fractionation | 3 |
| | Co-localization | 3 |
| | Co-purification | 4 |
| | PCA | 1 |
| | Reconstituted Complex | 30 |
| | Two-hybrid | 5 |
| *Macaca mulatta* | Affinity Capture-Western | 1 |
| *Homo sapiens* | Affinity Capture-Luminescence | 315 |
| | Affinity Capture-MS | 21501 |
| | Affinity Capture-RNA | 477 |
| | Affinity Capture-Western | 31023 |
| | Biochemical Activity | 6148 |
| | Co-crystal Structure | 845 |
| | Co-fractionation | 1038 |
| | Co-localization | 1645 |
| | Co-purification | 1263 |
| | Far Western | 602 |
| | FRET | 604 |
| | PCA | 513 |
| | Protein-peptide | 1522 |
| | Protein-RNA | 235 |

Continued on next page

Table S1: **Low throughput data.** The low throughput (< 200 interactions) datasets extracted from V164 of the BioGRID database. Interaction data were limited to physical interactions from eukaryotic species then split into individual datasets by study and species. Low throughput data were then grouped into datasets by BioGRID experimental type.

| **Species** | **Type** | **Interactions** |
|---|---|---|
| | Proximity Label-MS | 660 |
| | Reconstituted Complex | 17913 |
| | Two-hybrid | 10220 |
| *Canis lupus familiaris* | Affinity Capture-MS | 1 |
| | Affinity Capture-Western | 16 |
| | Co-localization | 2 |
| | Reconstituted Complex | 3 |
| *Sus scrofa* | Affinity Capture-MS | 32 |
| | Affinity Capture-Western | 12 |
| | Co-crystal Structure | 1 |
| | Reconstituted Complex | 3 |
| | Two-hybrid | 2 |
| *Bos taurus* | Affinity Capture-MS | 3 |
| | Affinity Capture-Western | 55 |
| | Biochemical Activity | 9 |
| | Co-crystal Structure | 2 |
| | Co-fractionation | 1 |
| | Far Western | 2 |
| | Reconstituted Complex | 17 |
| | Two-hybrid | 3 |
| *Ovis aries* | Affinity Capture-Western | 1 |
| *Oryctolagus cuniculus* | Affinity Capture-Western | 13 |
| | Biochemical Activity | 2 |
| | PCA | 1 |
| | Reconstituted Complex | 17 |
| *Cricetulus griseus* | Affinity Capture-Western | 1 |
| *Mus musculus* | Affinity Capture-Luminescence | 19 |
| | Affinity Capture-MS | 3469 |
| | Affinity Capture-RNA | 29 |
| | Affinity Capture-Western | 4807 |
| | Biochemical Activity | 372 |
| | Co-crystal Structure | 36 |
| | Co-fractionation | 195 |
| | Co-localization | 151 |
| | Co-purification | 41 |
| | Far Western | 64 |

Continued on next page

Table S1: **Low throughput data.** The low throughput (< 200 interactions) datasets extracted from V164 of the BioGRID database. Interaction data were limited to physical interactions from eukaryotic species then split into individual datasets by study and species. Low throughput data were then grouped into datasets by BioGRID experimental type.

| **Species** | **Type** | **Interactions** |
|---|---|---|
| | FRET | 33 |
| | PCA | 12 |
| | Protein-peptide | 151 |
| | Protein-RNA | 7 |
| | Proximity Label-MS | 55 |
| | Reconstituted Complex | 929 |
| | Two-hybrid | 845 |
| *Rattus norvegicus* | Affinity Capture-Luminescence | 4 |
| | Affinity Capture-MS | 801 |
| | Affinity Capture-RNA | 3 |
| | Affinity Capture-Western | 1061 |
| | Biochemical Activity | 96 |
| | Co-crystal Structure | 3 |
| | Co-fractionation | 61 |
| | Co-localization | 59 |
| | Co-purification | 12 |
| | Far Western | 13 |
| | FRET | 11 |
| | PCA | 3 |
| | Protein-peptide | 5 |
| | Protein-RNA | 2 |
| | Reconstituted Complex | 286 |
| | Two-hybrid | 219 |
| *Cavia porcellus* | Affinity.Capture-Western | 1 |
| | Biochemical Activity | 1 |
| | Co-purification | 1 |
| | Reconstituted Complex | 1 |
| *Vitis vinifera* | FRET | 1 |
| *Plasmodium falciparum* 3D7 | Co-crystal Structure | 1 |
| | Reconstituted Complex | 1 |
| | Two-hybrid | 1 |
| *Oryza sativa* Japonica Group | Affinity Capture-Western | 2 |
| | Biochemical Activity | 1 |
| | FRET | 4 |
| | PCA | 2 |
| | Reconstituted Complex | 5 |

Continued on next page

Table S1: **Low throughput data.** The low throughput (< 200 interactions) datasets extracted from V164 of the BioGRID database. Interaction data were limited to physical interactions from eukaryotic species then split into individual datasets by study and species. Low throughput data were then grouped into datasets by BioGRID experimental type.

| **Species** | **Type** | **Interactions** |
|---|---|---|
| | Two-hybrid | 26 |
| *Chlorocebus sabaeus* | Affinity Capture-Western | 4 |
| *Selaginella moellendorffii* | Reconstituted Complex | 2 |
| | Two-hybrid | 4 |
| *Aspergillus nidulans* FGSC A4 | Affinity Capture-MS | 49 |
| | Affinity Capture-Western | 5 |
| | Biochemical Activity | 1 |
| | PCA | 8 |
| | Two-hybrid | 1 |
| *Candida albicans* SC5314 | Affinity Capture-MS | 400 |
| | Affinity Capture-Western | 122 |
| | Biochemical Activity | 24 |
| | Co-crystal Structure | 3 |
| | Co-localization | 3 |
| | Co-purification | 6 |
| | Far Western | 2 |
| | PCA | 4 |
| | Protein-peptide | 2 |
| | Reconstituted Complex | 14 |
| | Two-hybrid | 60 |
| | Affinity Capture-Western | 1 |
| *Schizosaccharomyces pombe* 972h- | Affinity Capture-Luminescence | 1 |
| | Affinity Capture-MS | 2514 |
| | Affinity Capture-RNA | 170 |
| | Affinity Capture-Western | 1364 |
| | Biochemical Activity | 240 |
| | Co-crystal Structure | 45 |
| | Co-fractionation | 52 |
| | Co-localization | 336 |
| | Co-purification | 204 |
| | Far Western | 27 |
| | FRET | 9 |
| | PCA | 8 |
| | Protein-peptide | 30 |
| | Protein-RNA | 9 |
| | Proximity Label-MS | 1 |

Continued on next page

Table S1: **Low throughput data.** The low throughput (< 200 interactions) datasets extracted from V164 of the BioGRID database. Interaction data were limited to physical interactions from eukaryotic species then split into individual datasets by study and species. Low throughput data were then grouped into datasets by BioGRID experimental type.

| **Species** | **Type** | **Interactions** |
|---|---|---|
| | Reconstituted Complex | 469 |
| | Two-hybrid | 854 |
| *Saccharomyces cerevisiae* | Affinity Capture-Luminescence | 34 |
| | Affinity Capture-MS | 12580 |
| | Affinity Capture-RNA | 734 |
| | Affinity Capture-Western | 9993 |
| | Biochemical Activity | 1892 |
| | Co-crystal Structure | 590 |
| | Co-fractionation | 878 |
| | Co-localization | 695 |
| | Co-purification | 1875 |
| | Far Western | 86 |
| | FRET | 166 |
| | PCA | 1327 |
| | Protein-peptide | 279 |
| | Protein-RNA | 296 |
| | Proximity Label-MS | 81 |
| | Reconstituted Complex | 5021 |
| | Two-hybrid | 4909 |

Table S2: **High throughput data.** High throughput datasets extracted from V164 of the BioGRID database (>=200 interactions). Interaction data were limited to physical interactions (Ints) from eukaryotic species then split into individual datasets by species and by study based on PubMed ID (PMID).

| **Species** | **PMID** | **Ints** | **PMID** | **Ints** | **PMID** | **Ints** |
|---|---|---|---|---|---|---|
| *Arabidopsis thaliana* | 18535787 | 200 | 23449474 | 396 | 21094157 | 536 |
| | 24846764 | 222 | 21734647 | 423 | 21189294 | 555 |
| | 19429840 | 246 | 24823379 | 469 | 23166809 | 666 |
| | 15805477 | 255 | 20855607 | 482 | 20706207 | 811 |
| | 22065421 | 276 | 24556609 | 506 | 23667124 | 941 |
| | 21423366 | 346 | 22737156 | 529 | 21798944 | 5506 |
| | 20407024 | 395 | 21952135 | 534 | 24833385 | 12051 |
| *Caenorhabditis elegans* | 18692475 | 688 | 14704431 | 4307 | | |
| *Drosophila melanogaster* | 24114784 | 201 | 21447707 | 754 | 27626673 | 1800 |
| | 25453831 | 217 | 23749869 | 1217 | 22036573 | 10917 |
| | 22028469 | 386 | 15710747 | 1726 | 14605208 | 19886 |
| | 25242320 | 618 | 15575970 | 1759 | | |
| *Xenopus laevis* | 26673895 | 637 | | | | |
| *Homo sapiens* | 25192599 | 200 | 16196087 | 311 | 26687479 | 680 |
| | 24639526 | 201 | 21163940 | 311 | 15231748 | 692 |
| | 24244333 | 202 | 26777405 | 315 | 20562859 | 706 |
| | 25659154 | 203 | 23084401 | 317 | 17643375 | 709 |
| | 16159877 | 203 | 24797263 | 322 | 24255178 | 723 |
| | 15324660 | 208 | 24337577 | 323 | 16713569 | 764 |
| | 17314511 | 208 | 25147182 | 325 | 25281560 | 812 |
| | 22145905 | 208 | 23398456 | 326 | 20467437 | 853 |
| | 20186120 | 208 | 24344204 | 330 | 27621311 | 877 |
| | 19953087 | 211 | 25693804 | 331 | 22990118 | 947 |
| | 24705354 | 218 | 24623722 | 336 | 25963833 | 967 |
| | 24550385 | 219 | 22939624 | 363 | 23602568 | 984 |
| | 22493164 | 220 | 23824909 | 373 | 24332808 | 1008 |
| | 25756610 | 221 | 23752268 | 382 | 26673895 | 1037 |
| | 20020773 | 222 | 26167880 | 392 | 19738201 | 1068 |
| | 19295130 | 222 | 25324306 | 392 | 21516116 | 1125 |
| | 25277244 | 223 | 21319273 | 396 | 27880917 | 1321 |
| | 27637333 | 225 | 21903422 | 404 | 28611215 | 1488 |
| | 23349634 | 229 | 17620599 | 408 | 27432908 | 1510 |
| | 15175163 | 229 | 24189400 | 410 | 27499296 | 1514 |
| | 19060904 | 233 | 24999758 | 424 | 23443559 | 1514 |
| | 25515538 | 235 | 15231747 | 432 | 19615732 | 1538 |
| | 28378594 | 235 | 24366813 | 453 | 19322201 | 1669 |
| | 26641092 | 238 | 25036637 | 457 | 24711643 | 1736 |

Continued on next page

Table S2: **High throughput data.** High throughput datasets extracted from V164 of the BioGRID database (>=200 interactions). Interaction data were limited to physical interactions (Ints) from eukaryotic species then split into individual datasets by species and by study based on PubMed ID (PMID).

| Species | PMID | Ints | PMID | Ints | PMID | Ints |
|---|---|---|---|---|---|---|
| | 23125841 | 240 | 23463506 | 460 | 24457600 | 1776 |
| | 18782753 | 241 | 23455922 | 484 | 25609649 | 1822 |
| | 23246001 | 241 | 22623428 | 489 | 21832049 | 1893 |
| | 28190767 | 243 | 26831064 | 492 | 25921289 | 1921 |
| | 21182205 | 244 | 24722188 | 496 | 29128334 | 1975 |
| | 19156129 | 245 | 24778252 | 500 | 17353931 | 2050 |
| | 27926873 | 248 | 28561026 | 511 | 29117863 | 2296 |
| | 21081666 | 257 | 23455924 | 519 | 16169070 | 2449 |
| | 25737280 | 263 | 25468996 | 555 | 22658674 | 2473 |
| | 17043677 | 264 | 25241761 | 559 | 21900206 | 2604 |
| | 15604093 | 266 | 27591049 | 561 | 16189514 | 2605 |
| | 25315684 | 271 | 19549727 | 572 | 21988832 | 3433 |
| | 25900982 | 276 | 25640309 | 599 | 21145461 | 4540 |
| | 26514267 | 277 | 22365833 | 600 | 27173435 | 4905 |
| | 23667531 | 281 | 21044950 | 609 | 26638075 | 5347 |
| | 24163370 | 283 | 22268729 | 613 | 28431233 | 6219 |
| | 26389662 | 284 | 25814554 | 627 | 22863883 | 7129 |
| | 23383273 | 287 | 19380743 | 631 | 25416956 | 13424 |
| | 20811636 | 292 | 20936779 | 633 | 22939629 | 13917 |
| | 22119785 | 297 | 26618866 | 639 | 26496610 | 15215 |
| | 16055720 | 298 | 25852190 | 654 | 26344197 | 16795 |
| | 20133760 | 299 | 22586326 | 656 | 26186194 | 23709 |
| | 21244100 | 304 | 28065597 | 658 | 28514442 | 56467 |
| | 19690564 | 310 | 20211142 | 660 | | |
| *Mus musculus* | 16944949 | 244 | 28190767 | 375 | 24457600 | 1163 |
| | 25737280 | 258 | 27507650 | 435 | 28378742 | 1664 |
| | 20362541 | 274 | 29459677 | 462 | 28671696 | 1994 |
| | 22922362 | 360 | 21565611 | 617 | | |
| | 28706196 | 362 | 20211142 | 959 | | |
| *Rattus norvegicus* | 26316108 | 214 | 25931508 | 354 | 22871113 | 729 |
| *Plasmodium falciparum* 3D7 | 16267556 | 2435 | | | | |
| *Schizosaccharomyces pombe* 972h- | 26167880 | 201 | 24013502 | 225 | 24713849 | 931 |
| | 22119525 | 202 | 22540037 | 298 | 26771498 | 1939 |
| | 21652630 | 225 | 21386897 | 347 | | |
| *Saccharomyces cerevisiae* | 26804907 | 204 | 26217697 | 339 | 10688190 | 823 |
| | 15766533 | 205 | 21277287 | 352 | 21232131 | 933 |
| | 23209026 | 206 | 17922018 | 362 | 21460040 | 991 |

Continued on next page

Table S2: **High throughput data.** High throughput datasets extracted from V164 of the BioGRID database (>=200 interactions). Interaction data were limited to physical interactions (Ints) from eukaryotic species then split into individual datasets by species and by study based on PubMed ID (PMID).

| Species | PMID | Ints | PMID | Ints | PMID | Ints |
|---|---|---|---|---|---|---|
| | 22028667 | 216 | 12052880 | 370 | 23409723 | 995 |
| | 24374311 | 221 | 21825077 | 371 | 26493364 | 1080 |
| | 28539364 | 223 | 23911324 | 385 | 18805955 | 1089 |
| | 26777405 | 228 | 25172955 | 416 | 21734642 | 1122 |
| | 11743162 | 231 | 12150911 | 456 | 18757749 | 1206 |
| | 21926967 | 240 | 23712011 | 456 | 26668354 | 1244 |
| | 27550809 | 243 | 14660704 | 477 | 22271760 | 1320 |
| | 21944719 | 248 | 21107429 | 479 | 22940862 | 1412 |
| | 27905431 | 250 | 16606443 | 497 | 18719252 | 1610 |
| | 29361028 | 257 | 25792736 | 506 | 27863241 | 1686 |
| | 10900456 | 258 | 25228766 | 511 | 26524240 | 1751 |
| | 19106085 | 258 | 18087042 | 514 | 20489023 | 1849 |
| | 26392412 | 269 | 28096404 | 521 | 28768715 | 1875 |
| | 24500206 | 270 | 27552055 | 528 | 23222640 | 1886 |
| | 24152732 | 272 | 21558325 | 549 | 16093310 | 1941 |
| | 28325876 | 273 | 17634282 | 576 | 23332755 | 1973 |
| | 18046405 | 283 | 26849847 | 617 | 22199229 | 2007 |
| | 29738528 | 287 | 14759368 | 627 | 27821408 | 2053 |
| | 20595233 | 287 | 20170199 | 679 | 18467557 | 2395 |
| | 23945946 | 296 | 26831523 | 700 | 19840948 | 2541 |
| | 28539363 | 300 | 19536198 | 710 | 22875988 | 3207 |
| | 21070971 | 300 | 26673895 | 718 | 11805826 | 3210 |
| | 23403034 | 302 | 11283351 | 733 | 11805837 | 3598 |
| | 23831759 | 305 | 23904270 | 739 | 16319894 | 4021 |
| | 18667435 | 317 | 19841731 | 748 | 29158339 | 6390 |
| | 21386897 | 321 | 20508643 | 751 | 16429126 | 6531 |
| | 27373154 | 336 | 27580715 | 805 | 16554755 | 7075 |
| | 25452130 | 339 | 25026036 | 810 | | |

Table S3: **Genes of interest**. The protein components of signalling pathways, transcription factors and multipotency germline/stem-cell-related genes used to extract subnetworks in main text section 3.4.

| **Pathway** | **Name** | **Gene model** |
|---|---|---|
| Wnt/$\beta$-catenin canonical pathway | Wnt1 | HmN_000328000 |
| | Wnt2 | HmN_000112200 |
| | Wnt4 | HmN_000808600 |
| | Wnt5 | HmN_000108100 |
| | Wnt11a | HmN_002147900 |
| | Wnt11b | HmN_000022800 |
| | Fz5/8-FzA | HmN_000386300 |
| | Fz4-FzB | HmN_000319700 |
| | FzC | HmN_000494400 |
| | Fz1/2/3/6/7-FzD | HmN_000227100 |
| | FzE | HmN_000595200 |
| | Dishevelled-A | HmN_000446600 |
| | Dishevelled-B | HmN_000723000 |
| | GSK3 | HmN_000472700 |
| | APC | HmN_000462000 |
| | Axin | HmN_000572400 |
| | $\beta$-catenin A | HmN_000161700 |
| | $\beta$-catenin B | HmN_000192700 |
| | LEF1/TCF | HmN_000448500 |
| | WIF | HmN_000932100 |
| | SFRP | HmN_000556500 |
| | SFL | HmN_000359400 |
| Ca2+ - dependent Wnt pathway | Phospholipase C | HmN_000108000 |
| | Protein Kinase C | HmN_000418200 |
| | Calcineurin | HmN_000416800 |
| | CaMKII | HmN_000401300 |
| Planar cell polarity Wnt pathway | Strabismus | HmN_000676800 |
| | Prickle | HmN_000001700 |
| | RhoA | HmN_000173500 |
| | Rac | HmN_000316900 |
| | ROCK (Rho-Kinase) | HmN_000273400 |
| | JNK (JUN-Kinase) | HmN_000201000 |
| Notch pathway | Notch1 | HmN_000653600 |
| | Notch2 | HmN_000853800 |
| | Numb | HmN_000511800 |
| | HES1 | HmN_000974100 |
| | HES2 | HmN_000343800 |
| | HES3 | HmN_000973500 |

Continued on next page

Table S3: **Genes of interest**. The protein components of signalling pathways, transcription factors and multipotency germline/stem-cell-related genes used to extract subnetworks in main text section 3.4.

| Pathway | Name | Gene model |
|---|---|---|
| | Su(H) | HmN_000214100 |
| | Dl1 | HmN_000155700 |
| | Dl2 | HmN_000605500 |
| | Dl3 | HmN_000714400 |
| | Dl4 | HmN_000734400 |
| | Dl5 | HmN_000376400 |
| | Dl6 | HmN_000714300 |
| | Pres | HmN_000031500 |
| | Spdo | HmN_000234700 |
| | Ofut | HmN_000485500 |
| | Kuz | HmN_000907900 |
| | TACE | HmN_000264000 |
| | Nicastrin | HmN_000242800 |
| | Pen2 | HmN_000694400 |
| | APH1 | HmN_000691700 |
| | SMRT | HmN_000838900 |
| | Nle | HmN_000357200 |
| Hedgehog pathway | Cubitus Interruptus | HmN_000840400 |
| | Hedgehog | HmN_000068600 |
| | Disp | HmN_000570400 |
| | Patched | HmN_000602000 |
| | Smoothened | HmN_000930900 |
| | Su(fu) | HmN_000482400 |
| | Fu | HmN_000686000 |
| | Slimb | HmN_000834500 |
| | PKA1 | HmN_000251300 |
| | PKA2 | HmN_000254800 |
| | CK1-1 | HmN_003047480 |
| | CK1-3 | HmN_000690200 |
| | Caesin Kinase -CK1-3 | HmN_000690500 |
| | CK1-4 | HmN_000752200 |
| | CK1-5 | HmN_000297300 |
| | CK1-6 | HmN_000348900 |
| | GSK3-B1 | HmN_000354400 |
| | GSK3-B2 | HmN_000472700 |
| | CBP-5 | HmN_000756700 |
| | CBP-7 | HmN_000466400 |
| | Rsp-1 | HmN_000543900 |

Continued on next page

Table S3: **Genes of interest**. The protein components of signalling pathways, transcription factors and multipotency germline/stem-cell-related genes used to extract subnetworks in main text section 3.4.

| **Pathway** | **Name** | **Gene model** |
|---|---|---|
| | Igu | HmN_000604900 |
| | Rdx | HmN_000637800 |
| | Ttv | HmN_000215200 |
| | Sotv | HmN_000928900 |
| | Botv | HmN_001006500 |
| Hox transcription factors | Hox1 | HmN_000092500 |
| | Lox4/Abda | HmN_000400300 |
| | Hox4/Dfd | HmN_000772500 |
| | Lox5 | HmN_000187700 |
| | Post1a-1 | HmN_003005880 |
| | Post1a-2 | HmN_003005900 |
| | Post1b-1 | HmN_003005850 |
| | Post1b-2 | HmN_003005860 |
| | Post2 | HmN_000187500 |
| Stem-cell related | Ago1a | HmN_000114300 |
| | Ago1b | HmN_000950000 |
| | Ago2 | HmN_000658200 |
| | Bruno4 | HmN_000599500 |
| | DDX3 | HmN_000061900 |
| | DDXX | HmN_000797000 |
| | efTu a | HmN_000017000 |
| | efTu b | HmN_000074700 |
| | klf10 | HmN_000574300 |
| | klf5a | HmN_000013000 |
| | klf5b | HmN_000820000 |
| | MCM4 | HmN_000501500 |
| | MCM5 | HmN_000718100 |
| | Nanos | HmN_000753700 |
| | p63 | HmN_000797200 |
| | Pum1 | HmN_000758500 |
| | Pum2 | HmN_000516200 |
| | Pum3 | HmN_000125500 |
| | rbp4 | HmN_000255100 |
| | RNAhelME31B | HmN_000746800 |
| | Zfp1 | HmN_000668300 |
| | Zfp2 | HmN_000668400 |
| | Sox-PF1 | HmN_000208300 |
| | Sox-PF2 | HmN_000874900 |

Continued on next page

Table S3: **Genes of interest**. The protein components of signalling pathways, transcription factors and multipotency germline/stem-cell-related genes used to extract subnetworks in main text section 3.4.

| Pathway | Name | Gene model |
|---|---|---|
| | Sox-PF3 | HmN_000494900 |

Table S4: **Network clustering.** The proteins of the top ten clusters with in-cluster degree and MCODE protein score. Hub proteins (main text Table 2 are denoted *).

| Cluster | Degree | Score | Gene model | Description |
|---|---|---|---|---|
| Cluster 1 | 25 | 22.847 | HmN_000653200 | 40S ribosomal protein S12 |
| Cluster 1 | 26* | 23.778 | HmN_000490400 | 60S ribosomal protein L4 |
| Cluster 1 | 25 | 23.778 | HmN_000390700 | 60S ribosomal protein L7a |
| Cluster 1 | 26* | 23.778 | HmN_003026960 | 40S ribosomal protein S8 |
| Cluster 1 | 25 | 23.778 | HmN_000102000 | 60S ribosomal protein L21 |
| Cluster 1 | 26 | 23.778 | HmN_000101000 | 60S ribosomal protein L18a |
| Cluster 1 | 25 | 22.847 | HmN_000539200 | 60S ribosomal protein L19 |
| Cluster 1 | 25 | 22.847 | HmN_000257200 | 60S ribosomal protein L23a |
| Cluster 1 | 22 | 20.751 | HmN_000817200 | 40S ribosomal protein S7 |
| Cluster 1 | 26* | 23.778 | HmN_000077700 | 40S ribosomal protein S3a |
| Cluster 1 | 26* | 23.778 | HmN_000632900 | 40S ribosomal protein S23 |
| Cluster 1 | 25 | 22.847 | HmN_003033630 | 60S ribosomal protein L18 |
| Cluster 1 | 25 | 22.847 | HmN_003033620 | 60S ribosomal protein L18 |
| Cluster 1 | 26 | 23.778 | HmN_000384200 | 60S ribosomal protein L23 |
| Cluster 1 | 26 | 23.778 | HmN_003037990 | 60S ribosomal protein L37a |
| Cluster 1 | 26 | 23.778 | HmN_003028820 | 60S ribosomal protein L5 |
| Cluster 1 | 25 | 23.778 | HmN_000306800 | Proteasome subunit alpha type-3 |
| Cluster 1 | 26 | 23.778 | HmN_000211600 | 40S ribosomal protein S5 |
| Cluster 1 | 25 | 23.778 | HmN_000573200 | 60S ribosomal protein L24 |
| Cluster 1 | 26* | 23.778 | HmN_000002900 | 40S ribosomal protein S4 |
| Cluster 1 | 26 | 23.778 | HmN_000021200 | 60S ribosomal protein L7 |
| Cluster 1 | 26* | 23.778 | HmN_000899300 | 40S ribosomal protein S13 |
| Cluster 1 | 26 | 23.778 | HmN_000753600 | 40S ribosomal protein S11 |
| Cluster 1 | 26* | 23.778 | HmN_000932000 | 40S ribosomal protein S16 |
| Cluster 1 | 26 | 23.778 | HmN_000646200 | 40S ribosomal protein S19 |
| Cluster 1 | 25 | 22.847 | HmN_000538600 | 60S ribosomal protein L19 |
| Cluster 1 | 26* | 23.778 | HmN_000217500 | 40S ribosomal protein S2 |
| Cluster 2 | 9 | 7.644 | HmN_000759100 | DNA-directed RNA polymerase II subunit RPB11 |
| Cluster 2 | 12* | 8.077 | HmN_003006230 | DNA-directed RNA polymerase II subunit RPB1 |
| Cluster 2 | 11 | 7.152 | HmN_000249300 | DNA-directed RNA polymerases I, II, and III |
| Cluster 2 | 9 | 7.644 | HmN_000692800 | DNA-directed RNA polymerases I, II, and III |
| Cluster 2 | 11 | 7.152 | HmN_000212300 | DNA-directed RNA polymerases I, II, and III |
| Cluster 2 | 11 | 7.152 | HmN_003041820 | DNA-directed RNA polymerases I, II, and III |
| Cluster 2 | 9 | 7.644 | HmN_000692700 | DNA-directed RNA polymerases I, II, and III |
| Cluster 2 | 12 | 8.077 | HmN_003043630 | DNA-directed RNA polymerase II subunit RPB2 |
| Cluster 2 | 9 | 7.644 | HmN_000871800 | DNA-directed RNA polymerase II subunit RPB9 |
| Cluster 2 | 11 | 7.152 | HmN_000254300 | DNA-directed RNA polymerase II subunit RPB4 |
| Cluster 2 | 12 | 8.077 | HmN_000095200 | Mediator of RNA polymerase II transcription subunit 10 |

Continued on next page

Table S4: **Network clustering.** The proteins of the top ten clusters with in-cluster degree and MCODE protein score. Hub proteins (main text Table 2 are denoted *).

| Cluster | Degree | Score | Gene model | Description |
|---|---|---|---|---|
| Cluster 2 | 12 | 8.077 | HmN_000546000 | DNA-directed RNA polymerase II subunit RPB7 |
| Cluster 2 | 12 | 8.077 | HmN_000263100 | DNA directed RNA polymerase II subunit RPB3 |
| Cluster 3 | 7 | 6.000 | HmN_000645700 | Proteasome subunit beta type-2 |
| Cluster 3 | 7 | 6.000 | HmN_000417500 | Proteasome subunit beta type-7 |
| Cluster 3 | 7 | 6.000 | HmN_003004800 | Proteasome subunit alpha type-4 |
| Cluster 3 | 7 | 6.000 | HmN_000916400 | Proteasome subunit beta type-3 |
| Cluster 3 | 7 | 6.000 | HmN_000681300 | Proteasome subunit alpha type-2 |
| Cluster 3 | 6 | 6.000 | HmN_000813600 | 26S proteasome regulatory subunit 6B |
| Cluster 3 | 5 | 5.000 | HmN_000969400 | Proteasome maturation protein |
| Cluster 3 | 6 | 6.000 | HmN_003000770 | SWI/SNF-related matrix-associated actin-dependent regulator of chromatin subfamily A-like protein 1 |
| Cluster 4 | 4 | 7.000 | HmN_000362900 | Pre-mRNA-splicing factor ISY1 |
| Cluster 4 | 5 | 6.806 | HmN_000566300 | Pre-mRNA-splicing factor RBM22 |
| Cluster 4 | 5 | 6.806 | HmN_000845900 | U5 small nuclear ribonucleoprotein 40 kDa protein |
| Cluster 4 | 8 | 7.418 | HmN_000677000 | Pre-mRNA-splicing factor SYF1 |
| Cluster 4 | 6 | 6.533 | HmN_000719900 | Protein BUD31 homolog |
| Cluster 4 | 8 | 8.000 | HmN_003036700 | U5 small nuclear ribonucleoprotein 200 kDa helicase |
| Cluster 4 | 10* | 7.418 | HmN_002028700 | U2 small nuclear ribonucleoprotein A' |
| Cluster 4 | 5 | 8.000 | HmN_000463400 | SNW domain-containing protein 1 |
| Cluster 4 | 4 | 7.000 | HmN_000879100 | Small nuclear ribonucleoprotein E |
| Cluster 4 | 7* | 8.000 | HmN_000755100 | Microfibrillar-associated protein 1 |
| Cluster 4 | 10* | 7.418 | HmN_000632800 | 116 kDa U5 small nuclear ribonucleoprotein component |
| Cluster 5 | 5 | 4.691 | HmN_003049000 | Tubulin alpha-1B chain |
| Cluster 5 | 5 | 4.691 | HmN_003048990 | Tubulin alpha-1C chain |
| Cluster 5 | 8 | 4.667 | HmN_000747100 | Dynactin subunit 1 |
| Cluster 5 | 8 | 4.022 | HmN_000838500 | Dystonin |
| Cluster 5 | 5 | 4.691 | HmN_003038390 | Tubulin alpha-1C chain |
| Cluster 5 | 5 | 4.691 | HmN_003038410 | Tubulin alpha-1C chain |
| Cluster 5 | 8 | 4.022 | HmN_000742700 | n/a |
| Cluster 5 | 5 | 4.691 | HmN_003042330 | Tubulin alpha-1C chain |
| Cluster 5 | 5 | 4.691 | HmN_003038420 | Tubulin alpha-1B chain |
| Cluster 5 | 8 | 4.022 | HmN_000742800 | n/a |
| Cluster 5 | 5 | 4.691 | HmN_003038400 | Tubulin alpha-1C chain |
| Cluster 5 | 8 | 4.022 | HmN_000811700 | ADP-ribosylation factor-like protein 2 |
| Cluster 5 | 5 | 4.691 | HmN_003049010 | Tubulin alpha-1C chain |
| Cluster 6 | 5 | 5.000 | HmN_000175400 | Eukaryotic translation initiation factor 3 subunit I |
| Cluster 6 | 7 | 5.000 | HmN_000616600 | Eukaryotic translation initiation factor 3 subunit M |
| Cluster 6 | 3 | 5.067 | HmN_003005280 | COP9 signalosome complex subunit 5 |

Continued on next page

Table S4: **Network clustering.** The proteins of the top ten clusters with in-cluster degree and MCODE protein score. Hub proteins (main text Table 2 are denoted *).

| **Cluster** | **Degree** | **Score** | **Gene model** | **Description** |
|---|---|---|---|---|
| Cluster 6 | 6 | 5.000 | HmN_003044180 | Eukaryotic translation initiation factor 3 subunit E |
| Cluster 6 | 5 | 5.000 | HmN_000681800 | Eukaryotic translation initiation factor 3 subunit D |
| Cluster 6 | 3 | 5.727 | HmN_000167100 | Cop9 signalosome complex subunit |
| Cluster 6 | 5 | 5.000 | HmN_000273900 | Eukaryotic translation initiation factor 3 subunit G |
| Cluster 6 | 3 | 5.067 | HmN_000798600 | COP9 signalosome complex subunit 2 |
| Cluster 6 | 5 | 5.000 | HmN_000665900 | Eukaryotic translation initiation factor 3 subunit K |
| Cluster 7 | 4 | 4.000 | HmN_000272300 | U6 snRNA-associated Sm-like protein LSm5 |
| Cluster 7 | 4 | 4.000 | HmN_000888600 | Probable small nuclear ribonucleoprotein G |
| Cluster 7 | 4 | 4.000 | HmN_000138700 | Probable U6 snRNA-associated Sm-like protein LSm4 |
| Cluster 7 | 4 | 4.000 | HmN_000842400 | U6 snRNA-associated Sm-like protein LSm6 |
| Cluster 7 | 4 | 4.000 | HmN_000254100 | U6 snRNA-associated Sm-like protein LSm3 |
| Cluster 8 | 4 | 0.857 | HmN_000070500 | 14-3-3 protein epsilon |
| Cluster 8 | 4 | 0.857 | HmN_000209900 | 14-3-3 protein zeta/delta |
| Cluster 8 | 4 | 0.857 | HmN_000780400 | 14-3-3 protein sigma |
| Cluster 8 | 4 | 0.857 | HmN_000659700 | 14-3-3 protein beta:alpha |
| Cluster 8 | 5 | 1.048 | HmN_003037390 | Protein argonaute-2 |
| Cluster 8 | 5 | 1.048 | HmN_000551100 | TBC domain-containing protein kinase-like protein |
| Cluster 8 | 5 | 1.048 | HmN_000196600 | Serine/threonine-protein kinase PknB |
| Cluster 8 | 4 | 0.857 | HmN_000379400 | 14-3-3 protein zeta:delta |
| Cluster 8 | 5 | 1.048 | HmN_000735000 | Mitogen-activated protein kinase kinase kinase 1 |
| Cluster 9 | 5 | 0.250 | HmN_000309500 | Cell division control protein 42 homolog |
| Cluster 9 | 5 | 0.222 | HmN_000316900 | Ras-related C3 botulinum toxin substrate 1 |
| Cluster 9 | 4 | 0.222 | HmN_000153000 | Serine/threonine-protein kinase PAK 3 |
| Cluster 9 | 5 | 0.250 | HmN_003049740 | Cell division control protein 42 homolog |
| Cluster 9 | 4 | 0.222 | HmN_000447150 | Serine/threonine-protein kinase PAK 3 |
| Cluster 9 | 4 | 0.222 | HmN_000059800 | serine/threonine protein kinase PAK |
| Cluster 9 | 5 | 0.250 | HmN_000337200 | Cell division control protein 42 homolog |
| Cluster 9 | 4 | 0.200 | HmN_000493200 | serine/threonine protein kinase PAK |
| Cluster 9 | 4 | 0.222 | HmN_000261000 | Serine/threonine-protein kinase PAK 3 |
| Cluster 10 | 5 | 0.222 | HmN_000426100 | Serine/threonine-protein kinase N1 |
| Cluster 10 | 5 | 0.200 | HmN_003040810 | Protein kinase C beta type |
| Cluster 10 | 4 | 0.200 | HmN_000173500 | Transforming protein RhoA |
| Cluster 10 | 5 | 0.250 | HmN_000671200 | RAC-alpha serine/threonine-protein kinase |
| Cluster 10 | 5 | 0.222 | HmN_000849600 | Serine/threonine-protein kinase N2 |
| Cluster 10 | 4 | 0.222 | HmN_000530900 | Receptor of activated protein C kinase 1 |
| Cluster 10 | 4 | 0.200 | HmN_000788000 | Transforming protein RhoA |
| Cluster 10 | 4 | 0.200 | HmN_000537500 | Transforming protein RhoA |
| Cluster 10 | 4 | 0.250 | HmN_003004910 | Ras-related C3 botulinum toxin substrate 1 |

Table S5: **Differential expression.** The 176 differentially-expressed genes identified between adults and 5-day old, metamorphosing larvae that form a sub-network of 668 interactions (main text Figure 8).

| Gene model | Degree | log2FoldChange | padj | Description |
|---|---|---|---|---|
| HmN_003048980 | 8 | -12.84 | 3.14E-35 | Tubulin beta-1 chain |
| HmN_003017900 | 31 | -12.4 | 5.91E-234 | Tubulin beta chain |
| HmN_000757600 | 2 | -11.91 | 5.89E-29 | F-box/WD repeat-containing protein 7 |
| HmN_003000240 | 4 | -11.79 | 4.86E-35 | Tubulin alpha-3 chain |
| HmN_000221300 | 6 | -11.24 | 6.49E-45 | Tubulin alpha-4A chain |
| HmN_003049000 | 14 | -11.22 | 9.34E-218 | Tubulin alpha-1B chain |
| HmN_000300300 | 6 | -11.2 | 2.83E-27 | Casein kinase II subunit alpha |
| HmN_000994000 | 4 | -10.9 | 9.35E-26 | Ribosomal protein S6 kinase alpha-6 |
| HmN_000139600 | 24 | -9.65 | 2.63E-19 | RING-box protein 1 |
| HmN_000501400 | 16 | -9.57 | 1.57E-19 | E3 ubiquitin ligase complex SCF subunit sconC |
| HmN_003048860 | 24 | -9.36 | 2.04E-53 | Actin |
| HmN_003048990 | 12 | -9.23 | 0 | Tubulin alpha-1C chain |
| HmN_000483300 | 16 | -9.22 | 2.44E-21 | E3 ubiquitin ligase complex SCF subunit sconC |
| HmN_003044090 | 21 | -9.13 | 1.34E-32 | Tubulin beta-2B chain |
| HmN_003032460 | 6 | -8.89 | 1.23E-20 | Casein kinase II subunit alpha' |
| HmN_003036840 | 6 | -8.84 | 7.61E-17 | Casein kinase II subunit alpha' |
| HmN_003023870 | 6 | -8.37 | 5.56E-51 | Casein kinase II subunit alpha |
| HmN_003036830 | 6 | -8.29 | 2.13E-24 | Casein kinase II subunit alpha' |
| HmN_000063500 | 34 | -8.13 | 6.95E-36 | Cullin-1 |
| HmN_003006810 | 28 | -8.04 | 2.11E-42 | Histone acetyltransferase p300 |
| HmN_002187600 | 4 | -8.01 | 5.94E-14 | Actin |
| HmN_003035160 | 26 | -7.91 | 5.84E-222 | Tubulin beta-4B chain |
| HmN_000145000 | 5 | -7.73 | 5.17E-55 | Cell division control protein 2 homolog |
| HmN_000341700 | 1 | -7.45 | 6.68E-44 | mRNA decay activator protein ZFP36 |
| HmN_003049010 | 14 | -6.61 | 7.33E-82 | Tubulin alpha-1C chain |
| HmN_000259300 | 15 | -6.56 | 1.85E-92 | Serine/threonine-protein phosphatase PP1-gamma catalytic subunit |
| HmN_003038400 | 12 | -6.55 | 1.92E-36 | Tubulin alpha-1C chain |
| HmN_003003620 | 34 | -6.54 | 2.63E-117 | Cullin-1 |
| HmN_000145100 | 3 | -6.4 | 5.35E-117 | Cyclin-dependent kinase 9 |
| HmN_003035350 | 7 | -6.32 | 7.66E-07 | Heat shock cognate 71 kDa protein |
| HmN_000180200 | 3 | -5.91 | 2.50E-106 | Peroxiredoxin-1 |
| HmN_002145300 | 8 | -5.49 | 1.57E-40 | Histone-lysine N-methyltransferase SUV39H1 |
| HmN_003035310 | 7 | -5.25 | 1.05E-54 | Heat shock cognate 71 kDa protein |
| HmN_000341500 | 9 | -5.18 | 1.01E-67 | 14-3-3 protein gamma |
| HmN_000012700 | 6 | -5.15 | 4.24E-51 | General transcription factor IIE subunit 2 |
| HmN_000629900 | 34 | -4.93 | 2.87E-12 | Cullin-1 |
| HmN_003007120 | 14 | -4.77 | 9.63E-10 | DNA damage-binding protein 1 |

Continued on next page

Table S5: **Differential expression.** The 176 differentially-expressed genes identified between adults and 5-day old, metamorphosing larvae that form a sub-network of 668 interactions (main text Figure 8).

| Gene model | Degree | log2FoldChange | padj | Description |
|---|---|---|---|---|
| HmN_003012080 | 3 | -4.72 | 4.98E-60 | Peroxiredoxin-1 |
| HmN_000376000 | 7 | -4.62 | 3.49E-113 | Tubulin alpha c chain |
| HmN_003018090 | 11 | -4.58 | 1.88E-37 | NEDD8-conjugating enzyme Ubc12 |
| HmN_003018040 | 11 | -4.52 | 4.50E-43 | NEDD8-conjugating enzyme Ubc12 |
| HmN_003038410 | 12 | -4.47 | 1.10E-12 | Tubulin alpha-1C chain |
| HmN_000742800 | 20 | -4.12 | 8.95E-203 | n/a |
| HmN_000135100 | 4 | -4 | 3.55E-70 | Actin |
| HmN_003035360 | 7 | -3.68 | 1.03E-121 | Heat shock cognate 71 kDa protein |
| HmN_000398500 | 2 | -3.56 | 7.08E-283 | Collagen alpha-2(I) chain |
| HmN_000906000 | 4 | -3.41 | 2.54E-12 | Tyrosine-protein kinase Yes |
| HmN_000144900 | 33 | -3.1 | 2.09E-59 | Cyclin-dependent kinase 2 |
| HmN_000780400 | 25 | -3.1 | 2.71E-34 | 14-3-3 protein sigma |
| HmN_000768100 | 4 | -2.88 | 5.99E-83 | Actin |
| HmN_000826300 | 8 | -2.75 | 1.17E-22 | Probable ATP-dependent DNA helicase HFM1 |
| HmN_000472100 | 7 | -2.61 | 4.72E-52 | Tubulin alpha chain |
| HmN_000112700 | 1 | -2.41 | 1.46E-155 | NEDD8-conjugating enzyme UBC12 |
| HmN_000084600 | 2 | -2.33 | 3.22E-115 | Titin |
| HmN_003018120 | 10 | -2.17 | 4.83E-06 | NEDD8-conjugating enzyme Ubc12 |
| HmN_000695800 | 14 | -2.02 | 1.11E-19 | Ras-related C3 botulinum toxin substrate 1 |
| HmN_000214600 | 2 | -1.93 | 4.65E-123 | Glycogen [starch] synthase |
| HmN_003045210 | 6 | -1.86 | 6.92E-45 | Actin |
| HmN_003049740 | 7 | -1.84 | 7.73E-47 | Cell division control protein 42 homolog |
| HmN_000381000 | 22 | -1.62 | 3.17E-91 | Cullin-5 |
| HmN_000410600 | 2 | -1.6 | 1.47E-36 | E3 ubiquitin-protein ligase TRIM56 |
| HmN_003018190 | 10 | -1.59 | 1.75E-16 | NEDD8-conjugating enzyme Ubc12 |
| HmN_000037600 | 1 | -1.54 | 2.08E-35 | Ras related protein ral-a |
| HmN_003028770 | 29 | -1.5 | 9.78E-118 | Tubulin beta-1 chain |
| HmN_003028760 | 8 | -1.45 | 6.59E-21 | Tubulin beta-1 chain |
| HmN_000781500 | 2 | -1.42 | 2.19E-33 | Triple functional domain protein |
| HmN_002036900 | 14 | -1.24 | 9.72E-95 | Transgelin-3 |
| HmN_000490500 | 5 | -1.21 | 8.71E-24 | Ubiquitin-conjugating enzyme E2 H |
| HmN_000799800 | 52 | -1.14 | 2.21E-09 | Heat shock protein HSP 90-alpha |
| HmN_003032900 | 5 | -1.09 | 4.32E-47 | Dynamin-1 |
| HmN_000806400 | 6 | -1.04 | 5.95E-12 | Cyclin-dependent kinase 4 |
| HmN_003040810 | 9 | -1.02 | 8.19E-08 | Protein kinase C beta type |
| HmN_003041270 | 3 | 1.01 | 2.42E-45 | Tubulin beta-4B chain |
| HmN_000884600 | 2 | 1.01 | 7.15E-08 | SUMO-conjugating enzyme UBC9 |
| HmN_003004280 | 9 | 1.02 | 6.81E-98 | Tubulin alpha-1A chain |

Continued on next page

Table S5: **Differential expression.** The 176 differentially-expressed genes identified between adults and 5-day old, metamorphosing larvae that form a sub-network of 668 interactions (main text Figure 8).

| Gene model | Degree | log2FoldChange | padj | Description |
|---|---|---|---|---|
| HmN_000306800 | 32 | 1.03 | 3.82E-09 | Proteasome subunit alpha type-3 |
| HmN_000417500 | 7 | 1.04 | 3.11E-38 | Proteasome subunit beta type-7 |
| HmN_000566300 | 10 | 1.04 | 1.56E-28 | Pre-mRNA-splicing factor RBM22 |
| HmN_000569700 | 3 | 1.04 | 2.28E-16 | U2 small nuclear ribonucleoprotein B” |
| HmN_000493200 | 9 | 1.05 | 2.46E-12 | Serine/threonine protein kinase pak |
| HmN_000694000 | 6 | 1.07 | 2.01E-25 | Mitotic checkpoint protein BUB3 |
| HmN_000123500 | 1 | 1.09 | 5.34E-11 | Neurogenic differentiation factor 1 |
| HmN_000538000 | 10 | 1.11 | 1.14E-56 | 26S proteasome regulatory subunit 4 |
| HmN_003036340 | 31 | 1.12 | 2.65E-45 | Elongation factor 1-alpha 1 |
| HmN_000689500 | 3 | 1.12 | 4.15E-11 | Transcription initiation factor TFIID subunit 9 |
| HmN_000603600 | 2 | 1.15 | 6.14E-19 | Triple functional domain protein |
| HmN_003003820 | 39 | 1.18 | 3.57E-07 | Eukaryotic initiation factor 4A-III |
| HmN_000209900 | 29 | 1.19 | 1.59E-61 | 14-3-3 protein zeta/delta |
| HmN_000840000 | 13 | 1.19 | 3.75E-09 | Uncharacterized WD repeat-containing protein alr3466 |
| HmN_000407100 | 5 | 1.23 | 6.95E-41 | RNA helicase aquarius |
| HmN_000645700 | 7 | 1.24 | 8.32E-34 | Proteasome subunit beta type-2 |
| HmN_000806500 | 7 | 1.24 | 1.63E-26 | Cyclin-Y-like protein 1 |
| HmN_000109300 | 14 | 1.24 | 2.40E-25 | Cyclin-dependent kinase 7 |
| HmN_003028820 | 32 | 1.28 | 1.15E-107 | 60S ribosomal protein L5 |
| HmN_000612100 | 8 | 1.3 | 1.50E-78 | Small glutamine-rich tetratricopeptide repeat-containing protein alpha |
| HmN_000817200 | 27 | 1.31 | 1.47E-19 | 40S ribosomal protein S7 |
| HmN_000569800 | 3 | 1.36 | 1.30E-53 | U1 small nuclear ribonucleoprotein A |
| HmN_002028700 | 39 | 1.37 | 2.39E-53 | U2 small nuclear ribonucleoprotein A’ |
| HmN_000520400 | 17 | 1.39 | 8.98E-17 | Cyclin-dependent-like kinase 5 |
| HmN_000262700 | 15 | 1.42 | 1.59E-38 | Splicing factor 3A subunit 2 |
| HmN_000490400 | 33 | 1.43 | 5.98E-122 | 60S ribosomal protein L4 |
| HmN_000539200 | 32 | 1.44 | 1.28E-93 | 60S ribosomal protein L19 |
| HmN_003004800 | 15 | 1.52 | 3.83E-66 | Proteasome subunit alpha type-4 |
| HmN_003033620 | 29 | 1.52 | 3.89E-58 | 60S ribosomal protein L18 |
| HmN_000077200 | 4 | 1.55 | 3.90E-109 | E3 ubiquitin-protein ligase SIAH2 |
| HmN_000005800 | 5 | 1.57 | 6.48E-25 | p21-activated protein kinase-interacting protein 1 |
| HmN_000759700 | 3 | 1.58 | 1.26E-111 | Thioredoxin-dependent peroxide reductase |
| HmN_000052500 | 3 | 1.59 | 6.27E-11 | Cofilin |
| HmN_003028780 | 21 | 1.61 | 4.17E-111 | Tubulin beta-2B chain |
| HmN_000077700 | 41 | 1.66 | 2.78E-100 | 40S ribosomal protein S3a |
| HmN_002014200 | 11 | 1.68 | 1.09E-10 | F-box/WD repeat-containing protein 7 |
| HmN_000359200 | 6 | 1.7 | 7.53E-29 | Cyclin-dependent kinase 14 |

Continued on next page

Table S5: **Differential expression.** The 176 differentially-expressed genes identified between adults and 5-day old, metamorphosing larvae that form a sub-network of 668 interactions (main text Figure 8).

| Gene model | Degree | log2FoldChange | padj | Description |
|---|---|---|---|---|
| HmN_000002900 | 37 | 1.72 | 1.77E-180 | 40S ribosomal protein S4 |
| HmN_000390700 | 31 | 1.73 | 4.63E-143 | 60S ribosomal protein L7a |
| HmN_000600900 | 4 | 1.73 | 2.52E-16 | U6 snRNA-associated Sm-like protein LSm1 |
| HmN_000403100 | 2 | 1.75 | 1.42E-99 | Inosine-5'-monophosphate dehydrogenase 2 |
| HmN_000573200 | 29 | 1.75 | 3.43E-40 | 60S ribosomal protein L24 |
| HmN_000202800 | 6 | 1.77 | 9.67E-78 | Small nuclear ribonucleoprotein-associated protein N |
| HmN_000257200 | 26 | 1.77 | 4.70E-28 | 60S ribosomal protein L23a |
| HmN_000530900 | 8 | 1.79 | 1.32E-247 | Receptor of activated protein C kinase 1 |
| HmN_003038390 | 14 | 1.83 | 4.06E-129 | Tubulin alpha-1C chain |
| HmN_003026960 | 36 | 1.89 | 1.86E-113 | 40S ribosomal protein S8 |
| HmN_002125100 | 1 | 1.9 | 1.30E-76 | 40S ribosomal protein SA |
| HmN_000323900 | 10 | 1.93 | 8.08E-14 | Ubiquitin-like protein 4A-A |
| HmN_000248600 | 12 | 2.05 | 1.26E-92 | Mitogen-activated protein kinase 14 |
| HmN_000217500 | 41 | 2.24 | 4.40E-121 | 40S ribosomal protein S2 |
| HmN_000050800 | 13 | 2.25 | 3.14E-09 | Ubiquitin-conjugating enzyme E2 D1 |
| HmN_000320300 | 11 | 2.36 | 8.84E-14 | Peptidyl-prolyl cis-trans isomerase-like 1 |
| HmN_003037560 | 3 | 2.4 | 2.39E-08 | GTPase HRas |
| HmN_000062700 | 6 | 2.51 | 9.47E-11 | Cyclin-dependent kinase-like 1 |
| HmN_003044080 | 8 | 2.66 | 1.52E-101 | Tubulin beta-2B chain |
| HmN_000849500 | 3 | 2.71 | 5.58E-55 | E3 ubiquitin-protein ligase TRIM9 |
| HmN_000101000 | 29 | 2.79 | 2.09E-35 | 60S ribosomal protein L18a |
| HmN_003037570 | 3 | 2.82 | 9.14E-08 | GTPase HRas |
| HmN_000547500 | 1 | 2.87 | 2.37E-81 | Histone H2A |
| HmN_000052600 | 3 | 2.98 | 5.73E-91 | Actin-depolymerizing factor 1 |
| HmN_000604400 | 4 | 3.06 | 6.53E-288 | Late histone H1 |
| HmN_000899300 | 35 | 3.07 | 1.13E-43 | 40S ribosomal protein S13 |
| HmN_000950100 | 15 | 3.08 | 5.00E-58 | 60S ribosomal protein L27 |
| HmN_000578800 | 6 | 3.09 | 2.14E-40 | U6 snRNA-associated Sm-like protein LSm8 |
| HmN_000547600 | 12 | 3.18 | 3.07E-34 | Histone H2B |
| HmN_002003300 | 2 | 3.27 | 3.41E-158 | U1 small nuclear ribonucleoprotein C |
| HmN_000969400 | 5 | 3.3 | 8.52E-201 | Proteasome maturation protein |
| HmN_000336400 | 11 | 3.3 | 1.63E-42 | 40S ribosomal protein S15 |
| HmN_000545700 | 4 | 3.36 | 6.59E-86 | 40S ribosomal protein S30 |
| HmN_000888600 | 4 | 3.59 | 1.07E-23 | Probable small nuclear ribonucleoprotein G |
| HmN_000102000 | 28 | 3.66 | 3.59E-143 | 60S ribosomal protein L21 |
| HmN_000138700 | 6 | 3.78 | 2.08E-39 | Probable U6 snRNA-associated Sm-like protein LSm4 |
| HmN_000116700 | 5 | 3.83 | 3.08E-169 | Protein transport protein Sec61 subunit beta |
| HmN_000735000 | 16 | 3.84 | 1.10E-155 | Mitogen-activated protein kinase kinase kinase 1 |

Continued on next page

Table S5: **Differential expression.** The 176 differentially-expressed genes identified between adults and 5-day old, metamorphosing larvae that form a sub-network of 668 interactions (main text Figure 8).

| Gene model | Degree | log2FoldChange | padj | Description |
|---|---|---|---|---|
| HmN_000646200 | 31 | 3.84 | 2.23E-60 | 40S ribosomal protein S19 |
| HmN_000812800 | 10 | 3.86 | 4.56E-53 | Cyclin-dependent kinases regulatory subunit 2 |
| HmN_000396000 | 6 | 4.05 | 4.46E-125 | Small nuclear ribonucleoprotein Sm D2 |
| HmN_000932000 | 37 | 4.06 | 3.32E-253 | 40S ribosomal protein S16 |
| HmN_000632900 | 35 | 4.06 | 2.17E-117 | 40S ribosomal protein S23 |
| HmN_000356900 | 18 | 4.3 | 6.87E-57 | 60S ribosomal protein L27a |
| HmN_000163800 | 5 | 4.36 | 8.40E-80 | Activated RNA polymerase II transcriptional coactivator p15 |
| HmN_000384200 | 31 | 4.37 | 3.84E-50 | 60S ribosomal protein L23 |
| HmN_000102700 | 6 | 4.41 | 2.14E-100 | Small nuclear ribonucleoprotein Sm D1 |
| HmN_003035300 | 13 | 4.54 | 1.20E-222 | Heat shock cognate 71 kDa protein |
| HmN_000652600 | 6 | 4.59 | 2.11E-112 | Histone H2A |
| HmN_000068700 | 17 | 4.63 | 6.64E-163 | 60S acidic ribosomal protein P1 |
| HmN_000753600 | 31 | 4.84 | 1.32E-303 | 40S ribosomal protein S11 |
| HmN_000254100 | 7 | 5.05 | 2.86E-31 | U6 snRNA-associated Sm-like protein LSm3 |
| HmN_000653200 | 26 | 5.23 | 6.13E-121 | 40S ribosomal protein S12 |
| HmN_000391400 | 6 | 5.48 | 1.03E-90 | Small nuclear ribonucleoprotein Sm D3 |
| HmN_000824700 | 9 | 5.52 | 1.69E-60 | 40S ribosomal protein S28 |
| HmN_000842400 | 8 | 5.53 | 8.04E-23 | U6 snRNA-associated Sm-like protein LSm6 |
| HmN_002101000 | 17 | 5.54 | 4.78E-158 | 60S acidic ribosomal protein P1 |
| HmN_000879100 | 12 | 5.58 | 1.17E-60 | Small nuclear ribonucleoprotein E |
| HmN_000219600 | 15 | 5.66 | 3.33E-82 | 60S acidic ribosomal protein P2 |
| HmN_000272300 | 4 | 5.88 | 4.05E-13 | U6 snRNA-associated Sm-like protein LSm5 |
| HmN_000070400 | 10 | 6.28 | 8.12E-178 | 40S ribosomal protein S17 |
| HmN_000154900 | 7 | 6.31 | 2.18E-68 | Small ubiquitin-related modifier 2 |
| HmN_000236800 | 10 | 7.32 | 3.53E-86 | 40S ribosomal protein S21 |
| HmN_003037990 | 30 | 8.59 | 6.62E-35 | 60S ribosomal protein L37a |
